# Individual yeast cells signal at different levels but each with good precision

**DOI:** 10.1101/2020.01.15.906867

**Authors:** Steven S. Andrews, Roger Brent

## Abstract

Different isogenic cells have been shown to exhibit widely varying responses to the same extracellular signal. Based on the assumption that this variation arises from noise in the signaling pathways that cells use to transmit information from surface to nucleus, recent publications asserted that single cells cannot detect their surroundings accurately. Here, we analyze existing data on gene expression induced by the *Saccharomyces cerevisiae* pheromone response system, finding that individual cells signal consistently over time, implying that response variation arises primarily from stable cell-to-cell differences rather than signaling noise. Individual cells transmit at least 2.7 bits of information through the pheromone response system, enabling each cell to distinguish between at least 6 pheromone concentrations. In principle, cells can gain further precision by internally referencing these responses with measurements of constitutively expressed genes. Combination with prior results shows that only about 6% of total response variation arises from signaling pathway noise.

**One-sentence summary:** Single yeast cells signal consistently over time, indicating that their signaling pathways transmit information precisely.

## Main text

Cell signaling systems sense extracellular conditions and transmit information about them into the cell. The cell then uses the information to make decisions, such as whether to grow, undergo apoptosis, or differentiate. Incorrect decisions can lead to an undesirable fate, so one would reasonably expect that cell signaling systems would have evolved to transmit information accurately, as indeed was the assumption for many years (e.g. ref. [1]).

However, the actual amount of information transmitted through cell signaling systems was not been quantified until recently, when several groups of researchers applied chemical stimuli to isogenic collections of single cells and measured the responses [2–5]. They observed wide variation, which they quantified with information theory (see refs. [6–10]) to show that the channel capacity for a single cell, meaning the maximum information that can be discerned about the stimulus level based on one cell’s response, is only about one bit. This number corresponds to two states, implying that a cell’s response is adequate for determining whether a stimulus is present or not, but cannot give further detail about its concentration. This suggested that single cells cannot sense their surroundings precisely but may need to combine information from multiple sources to choose appropriate responses [2, 11, 12].

We propose a different interpretation, which is that the highly variable responses arise from temporally stable cell-to-cell differences instead of noisy biochemical signals. In this interpretation, which agrees with substantial prior work [13–16], individual cells behave differently from each other, but each is able to distinguish between different external conditions reasonably precisely by itself. By extension, individual cells might be able to make well-informed decisions on their own from relatively few inputs.

We tested this possibility by analyzing published data on signaling by the yeast (*Saccharomyces cerevisae*) pheromone response system (PRS). This system, shown in Figure 1A, is a prototypical G-protein signaling system, bears close homology to many mammalian signaling systems and has been studied thoroughly by us and others [17–21]. It transmits information about pheromone (α-factor) binding at cell surface receptors to the cell nucleus, where the signal then induces expression of several genes. The cells that we investigated expressed yellow fluorescent protein (YFP) from a pheromone-responsive promoter (P_PRM1_) and either red or cyan fluorescent protein (RFP or CFP) from the constitutive P_ACT1_ promoter [18, 19] (see Supplementary Materials section 1 (SM-1)).

**Figure 1.**
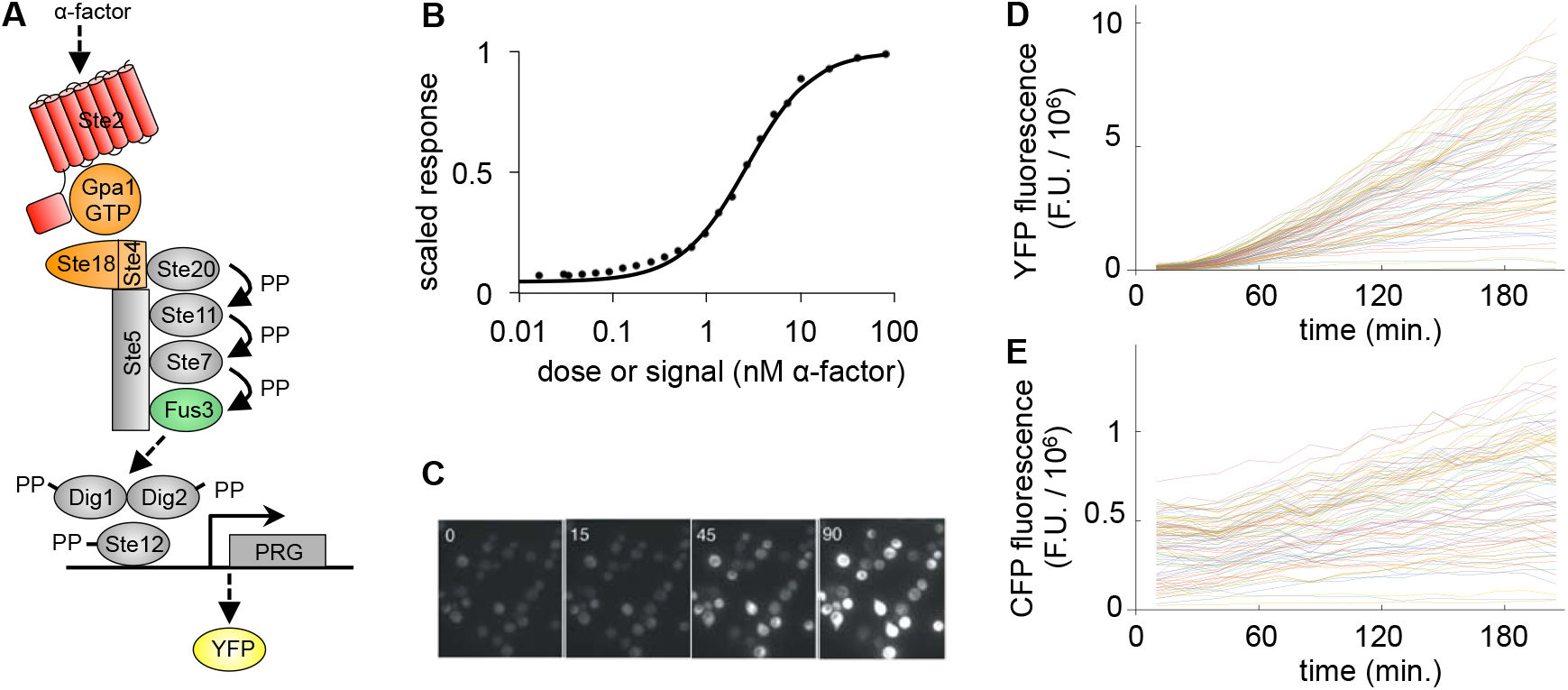
(A) The yeast PRS in mating type **a** cells. Pheromone (α-factor) binds to G-protein coupled receptors (Ste2), causing dissociation of heterotrimeric G-proteins (Gpa1/Ste18/Ste4), which recruits Ste5 scaffold proteins to the cell membrane and induces signaling through a MAP kinase cascade (Ste11, Ste7, Fus3, and Kss1); the MAP kinases activate Ste12 transcription factors which then promote expression of pheromone-responsive genes, in this case leading to YFP expression [17]. The pheromone concentration is the signal and YFP fluorescence is the response. (B) Dose-response curve for the PRS. Points represent experimental data from ref. [19] and the line represents a Hill function fit to the data from ref. [20]. Points and fit were scaled to approach a maximum of 1. (C) Microscope image of YFP fluorescence from several cells at 0, 15, 45, and 90 minutes after measurement began, showing temporally consistent variation. Copied with permission from ref. [18]. (D) YFP fluorescence from 90 individual cells over time after pheromone addition, quantified in arbitrary fluorescence units (F.U.). (E) CFP fluorescence, expressed from a constitutive promoter, from the same 90 individual cells. Data for panels D and E were measured for ref. [18] and made publically available in ref. [22].

We used two data sets in our analysis, from essentially identical cells. The first, from ref. [19] and shown in Figure 1B, represents the steady-state dose-response function for the PRS. It was collected using flow cytometry in which each cell’s response was computed as the ratio of pheromone-induced YFP expression to constitutive RFP expression in the same cell, which was then averaged over many cells. We required a dose-response function that was defined everywhere rather than only at discrete pheromone concentrations, so we used a Hill function fit to these experimental data (from ref. [20]) in our analysis rather than the raw data themselves (SM-2).

The second data set, described in refs. [18, 22], represents single-cell responses to pheromone stimulation. At each of 5 pheromone concentrations, these data report the YFP and CFP fluorescence intensities of roughly 100 individual yeast cells at 14 equally spaced time points after pheromone addition, measured by microscopy (Fig. 1C). We minimally filtered the data to remove entries for dead cells, badly segmented cells, and outlier measurements (SM-3). Figure 1D shows the YFP expression for 90 individual cells, each as a separate line, after stimulation by 20 nM pheromone. It shows that YFP concentrations stayed low initially and then increased nearly linearly over time for at least 3 hours. As neither pheromone nor intracellular YFP were degraded significantly during these experiments, the PRS within each cell must have signaled at a nearly constant rate. Figure 1E shows that the CFP concentration, from a constitutively expressed promoter in the same cells, did not respond to pheromone stimulation but only increased slightly due to cell growth.

We quantified signaling fidelity in two ways. The first, shown with yellow shading in Figure 2, treated all variation as signaling uncertainty, while ignoring the difference between cell-to-cell variation and within-cell signaling noise, to allow a meaningful comparison with prior results [2–5]. The second, shown with green shading, distinguished between these two types of variation so that we could quantify information transmission within single cells.

**Figure 2.**
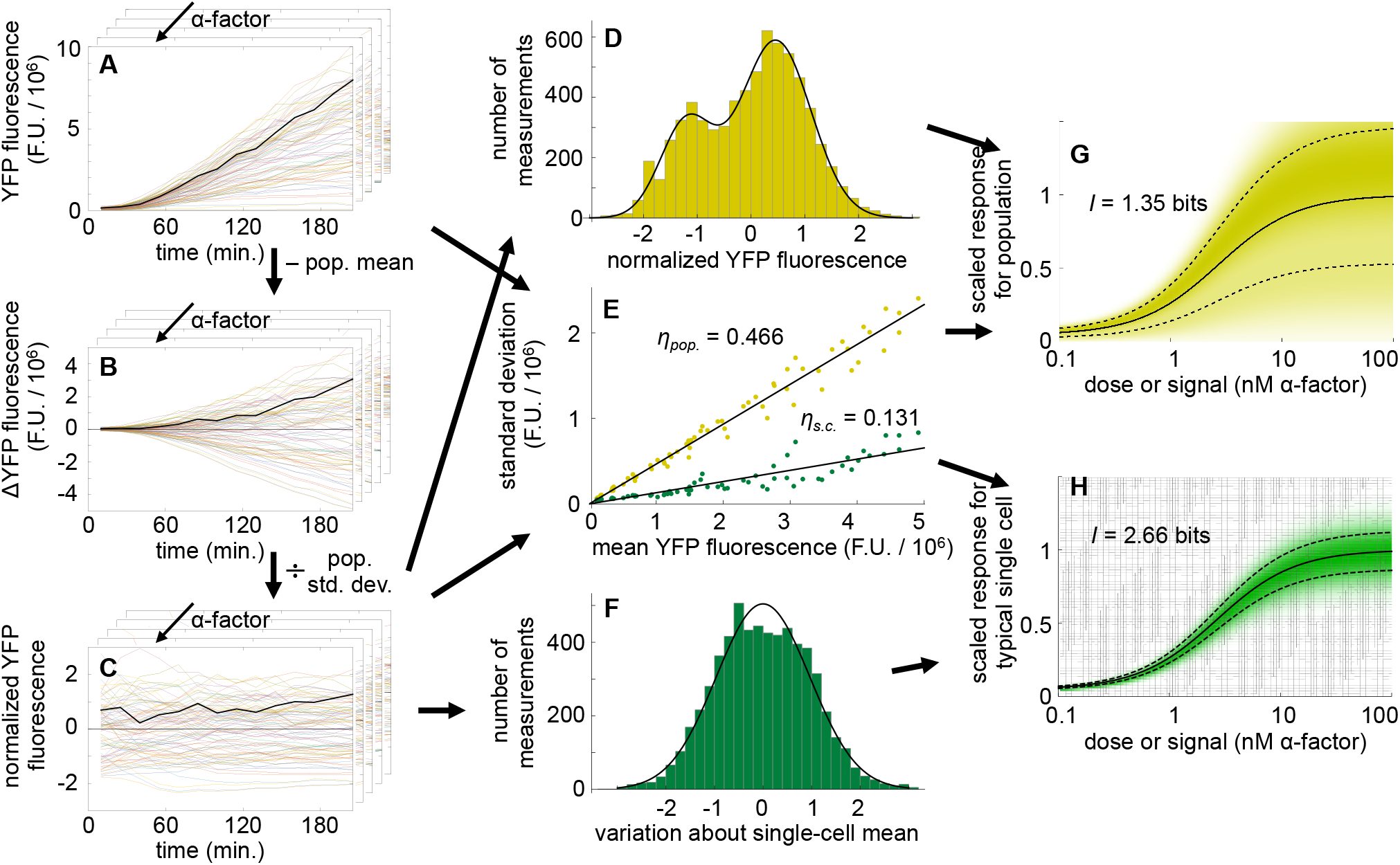
Channel capacity computation method. (A) Filtered single-cell YFP fluorescence data, where each line represents fluorescence from a single cell over time; a representative cell is shown in black. Each layer represents a different pheromone dose. (B) The same data, but with the population mean (all cells at a given time point and pheromone dose) fluorescence values subtracted; the same cell is shown in black. (C) The normalized single-cell data, in which fluorescence difference values were divided by the population standard deviation; the same cell is shown in black. (D) Distribution of normalized fluorescence values from panel C. The line is a best fit with a sum of two Gaussians. (E) Correlation between mean and standard deviation values for the cell population in yellow and an average single cell in green. Each point represents a single time point and pheromone concentration. Lines are best fits that were constrained to intersect the origin. (F) Distribution of normalized fluorescence values about the normalized single-cell mean values, scaled with the single-cell standard deviations. The line is a best fit with a single Gaussian. (G) Signal-response-variation (SRV) curve for the cell population. Shading represents the conditional response probability, *p*(*r*|*s*), the solid line represents the mean of this distribution, 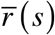, and the dashed line represents the standard deviation of this distribution. (H) SRV curve for an average single cell, showing lower variation and hence greater information transfer.

### Combined variation

From the single-cell data for YFP fluorescence (hereafter, just “single-cell data”; Fig. 2A), we computed the mean and standard deviation YFP fluorescence for the entire cell population at each time point and pheromone dose. A scatter plot of these data (Fig. 2E, yellow points) shows a linear relationship, implying a constant coefficient of variation (η_pop._ = 0.466) between the mean responses of the cell population and the variation across the population. This agrees with prior work that focused on high-abundance proteins and is consistent with the idea that variation is dominated by extrinsic noise, including cell-to-cell variation [3, 18, 23, 24] (intrinsic noise, such as protein expression that varies due to mRNA fluctuations, results in the coefficient of variation decreasing in proportion to the square root of the mean value).

We also normalized the single-cell data by subtracting the population mean at each time point (Fig. 2B) and then dividing by the population standard deviation at each time point (Fig. 2C). Each of these normalized fluorescence values represents the brightness of a particular cell relative to the population as a whole, at any given time point. These values exhibited a bimodal distribution (Fig. 2D), likely arising from cells in different growth phases.

Next, we computed what we call a signal-response-variation (SRV) curve (Fig. 2G; SM-4). Here, the mean response is the steady-state dose-response curve, the response standard deviation comes from the η_pop._ value of 0.466, and the shape of the response distribution at any given pheromone concentration is the bimodal distribution of YFP fluorescence values. Shading in the SRV curve represents the probability of observing response *r* given that the signal is *s*, which is called the conditional response probability and written as *p*(*r*|*s*). We computed the channel capacity from the conditional response that is shown in the SRV curve using the Blahut-Arimoto algorithm [25, 26] (SM-5), finding a result of 1.35 bits, which we call the population channel capacity. This means that knowledge of the YFP fluorescence from any randomly chosen single cell is sufficient to convey about 1.35 bits of information about the pheromone concentration. This corresponds to 2.5 different states (2^1.35^), so knowing the YFP fluorescence from one cell can generally indicate whether pheromone is present or absent but does not give much more detail than that. This agrees with the about 1 bit of channel capacity found in previous studies of other cells [2–5].

### Separated variation

To separate the effects of cell-to-cell variation from within-cell signaling noise, we assumed that the different slopes of the single-cell data traces shown in Figure 2A arose from temporally stable cell-to-cell variation and that the small “wiggles” within each of these traces represented signaling noise. In principle, these wiggle sizes could be computed by fitting smooth lines to the YFP fluorescence data and then computing residuals from them. However, such an approach would introduce artifacts from the necessarily imperfect fits to the non-linear time dependence. Thus, we chose a less direct but more neutral approach. Starting with the normalized data (Fig. 2C), we computed the difference between each normalized data point and the mean value for that cell over time to yield the normalized single-cell noise value for each data point. Then, at each time point and pheromone dose, we computed the root mean square (rms) average of these noise values over all cells. Finally, we removed the normalization by multiplying each rms average noise value by the population standard deviation. The result is an average of the noise amount (or wiggle size) over all cells, at each time point and pheromone dose, in units of YFP fluorescence. Figure 2E shows the results with green dots, plotting these single-cell standard deviations against the mean fluorescence values. It again shows a linear relationship between the mean and standard deviations, but now with a coefficient of variation of η_s.c._ = 0.131. Much as before, the linear relationship is consistent with variation being dominated by extrinsic noise rather than the intrinsic noise that arises from the stochastic birth and death processes of individual transcripts or proteins [24, 27].

We also computed the distribution of single-cell noise values about the respective single-cell mean values. To account for the fact that brighter cells exhibited more noise, we divided the normalized single-cell noise values by the normalized single-cell mean values to yield noise values that could be compared between different cells, as well as between different time points and pheromone doses. Figure 2F shows the distribution of these single-cell noise values. It agrees reasonably well with a Gaussian, likely indicating that single-cell noise values arise from a sum of many factors, such as biochemical stochasticity in different signaling proteins and various types of measurement error.

Combining these results created an SRV curve for a typical single cell (Fig. 2H; SM-6), meaning one whose response matches that of the population average and whose deviations arising from noise agree with the average deviations. In particular, this typical cell’s noise deviations are Gaussian distributed and have a coefficient of variation of η_s.c._. The single-cell channel capacity, computed from this SRV curve, is 2.66 bits. This value represents the amount of information that is transmitted from a typical cell’s pheromone receptors to its induced protein expression. It implies that a typical cell can signal precisely enough to distinguish between at least 6 different pheromone concentrations (2^2.66^). Because the data also included measurement noise, 2.66 bits is a lower limit and the true single-cell channel capacity may be substantially higher. This finding that cells can signal precisely is consistent with experimental results [28–31].

The difference between the two channel capacity results is akin to the formal difference between accuracy and precision. Accuracy represents the closeness of measurements to a true value whereas precision represents measurement reproducibility. A scientist who measured the pheromone concentration by using the YFP fluorescence from one yeast cell would get a fairly inaccurate result, conveying only 1.35 bits of information. However, if the scientist had previously calibrated this cell’s response using some known pheromone concentration, then a new measurement would convey 2.66 bits of information due to good signaling precision.

If a cell is only exposed to pheromone once in its life, then it would seem that its ability to transmit signals precisely would be wasted because it could still only estimate the external pheromone concentration to 1.35 bits. However, cells may be able to do better than this using the fact that expression levels of different genes tend to be correlated [18, 27, 32]. For example, one might imagine that a cell could compare the expression of a pheromone-responsive gene with that of a constitutively expressed gene, using the latter as an internal standard. We investigated this possibility by quantifying the information that a cell would learn if its YFP fluorescence value were divided by the simultaneously measured CFP value, computing the population and single-cell channel capacities as before. This normalization increased the population channel capacity to 2.01 bits, implying that a cell could accurately distinguish about 4 different pheromone concentrations with a single measurement. It also increased the single-cell channel capacity to 2.92 bits, enabling a cell to distinguish about 8 different pheromone concentrations. Such cellular comparison is not wholly unreasonable, given the fact that the PRS is already able to sense the fraction of receptors bound by ligand, as opposed to only the absolute number of bound receptors [6, 33]. By extension, cells could use multiple internal standards, and/or standards that correlate particularly closely with pheromone-responsive gene expression, to further improve information transmission. Cells could also improve signaling precision (but not accuracy) by measuring pheromone repeatedly to improve information content by signal averaging [15] (SM-8).

We wondered whether all cells signal similarly or if those that express more YFP might be able to signal more precisely than those that express less. To test this, we grouped cells into low, medium, and high brightness categories based on their normalized mean YFP fluorescence values (Fig. 2C). Repeating the single-cell channel capacity calculations showed that the channel capacity was slightly higher for cells in medium and high brightness groups, but not by much (Fig. 3). This shows that the vast majority of these cells signaled precisely, regardless of their gene expression capacity.

**Figure 3.**
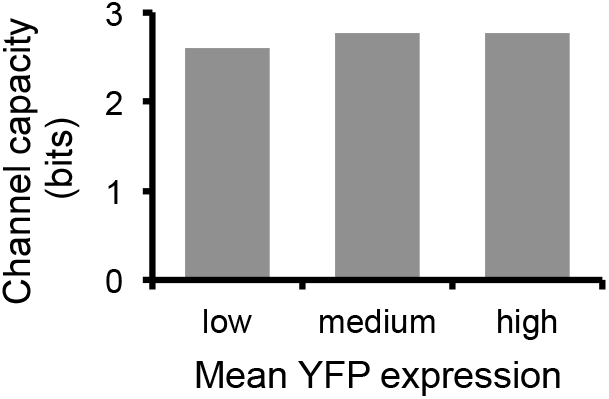
Channel capacity as a function of mean expression. We divided cells into low, medium, and high YFP expression groups based upon their normalized YFP fluorescence values and computed the single-cell channel capacity for each group of cells.

In previous work [18], we showed that total variance in pheromone-responsive expression, η^2^(*y*), can be expressed as a sum of four variances: η^2^(*L*) and η^2^(*G*) represent cell-to-cell variation in the capacity of the signaling pathway to transmit signals and in the capacity of the gene expression system to express genes, respectively, and η^2^(λ) and η^2^(γ) represent stochastic noise within the signaling pathway and in gene expression, respectively (an additional term accounts for correlations between pathway and expression capacity, but is often ignored, including here; see SM-7). In this formalism, our η_*pop.*_ value gets squared to give η^2^(*y*) = 0.217, which agrees well with prior results [18]. Also, our η_*s.c*._ value gets squared to give η^2^*s.c.* = 0.017. Dividing this by η^2^(*y*) shows that only about 8% of the total variation arises from noise within a cell, while 92% of variation arises from cell-to-cell differences. Furthermore, the single cell variance can be broken down as η^2^*s.c.* = η^2^(λ)+η^2^(γ), and the gene expression noise is about 2% of total variation [18], which shows that about 6% of total output variation arises from signaling pathway noise. Thus, again, we find that signaling pathway noise is a small contributor to the total response variation.

Based on the structural similarities between different eukaryotic signaling systems, along with the finding that the channel capacity for populations of cells is consistently about 1 bit, we suspect that our finding of precise signaling in single yeast cells applies to other signaling systems as well. If so, a wide variety of individual cells might be able to make well-informed decisions on their own.

## Acknowledgements

We thank Alejandro Colman-Lerner, Richard Yu, Gustavo Pesce, Alan Bush, and Bill Peria for assistance with the experimental data sets and helpful discussions. This work was funded by NCI R21 CA223901 to RB and a Simons Foundation Fellowship awarded to SA.

## Supplementary Information

### 1. Experimental data

#### 1.1. Experimental dose-response data

The data that we used for our steady-state dose-response curve was originally presented in Figure 2a of ref. (*1*), where it was shown with triangle symbols and described as “reporter gene expression output corrected for known sources of cell-to-cell variation (pathway output *P*)”. Richard Yu gave us the original data values for use in this and other work (see below and the supplementary information of ref. (*2*)). The original work did not describe some aspects of these data in detail, so we describe them here based upon information in ref. (*1*), the supplementary information for that paper, and personal communications with Richard Yu and Gustavo Pesce.

The data were collected on *S. cerevisiae* haploid strain RY2073 (not strain ACL379, as reported in ref. (*2*)). RY2073 was derived from TCY3154, which is from ACL379, which is from YAS245-5C, which is a W303 derivative. The RY2073 strain has the following genotype, from the supplementary information of ref. (*1*):

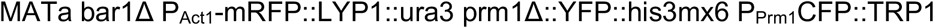

In more detail, RY2073 is mating type **a**, has the *BAR1* gene removed (to prevent pheromone degradation by Bar1 protein), has the *LYP1* gene removed (to confer resistance to the yeast poison thialysine), has the PAct1 promoter and *mRFP* gene inserted at the *LYP1* site (for constitutive expression of mRFP), has the *URA3* gene also inserted at the *LYP1* site (for selection on uracil-deficient plates), has the *PRM1* gene removed (a pheromone-responsive gene that codes for a protein that is involved in membrane fusion during mating), has the *YFP-His3MX6* gene inserted into the *PRM1* site and under control of the P_Prm1_ promoter (for pheromone-responsive YFP expression), and has a *CFP* gene that is expressed from another P_Prm1_ promoter and placed adjacent to the wild-type *TRP1* gene.

Fluorescence data were collected on these cells by sonicating exponentially growing cultures, diluting cells into fresh media, and then incubating them with appropriate concentrations of pheromone. Gene expression was stopped after 3 hours (for the steady-state signaling condition) by adding cycloheximide. RFP and YFP fluorescence was then measured on individual cells using high throughput flow cytometry. CFP fluorescence was not of interest and so was not intentionally excited and was blocked using optical filters.

These resulting experimental data were filtered to remove dead cells (about 2-6% of cells burst while trying to make a shmoo), which have a distinctive fluorescence pattern. The ratios of YFP to RFP fluorescence values for the remaining cells were used in the remainder of the analysis. The following table, also presented in ref. (*2*), lists these data values. The column labeled “<y>/<r>” represents the fluorescence value ratios and the column labeled “response” shows our re-scaled version of the raw data. It was adjusted so that the basal response would equal the experimental value when no pheromone was added and scaled to make the maximum of a Hill function that was fit to these data equal to 1 (see SM-2).

**Table SM-1.1.** Dose-response data for yeast PRS.

| $[\alpha]$ (nM) | $\langle y \rangle / \langle r \rangle$ | response | $[\alpha]$ (nM) | $\langle y \rangle / \langle r \rangle$ | response |
| --- | --- | --- | --- | --- | --- |
| 0 | 0.466 | 0.0470 | 0.00141 | 0.563 | 0.0571 |
| 1E-10 | 0.467 | 0.0471 | 0.00260 | 0.582 | 0.0590 |
| 1.84E-10 | 0.468 | 0.0472 | 0.00478 | 0.617 | 0.0627 |
| 3.39E-10 | 0.480 | 0.0485 | 0.00880 | 0.647 | 0.0658 |
| 6.23E-10 | 0.470 | 0.0475 | 0.0162 | 0.704 | 0.0716 |
| 1.15E-09 | 0.468 | 0.0473 | 0.0298 | 0.753 | 0.0767 |
| 2.11E-09 | 0.404 | 0.0406 | 0.0333 | 0.715 | 0.0728 |
| 3.88E-09 | 0.438 | 0.0441 | 0.0467 | 0.758 | 0.0772 |
| 7.14E-09 | 0.455 | 0.0459 | 0.0653 | 0.820 | 0.0837 |
| 1.31E-08 | 0.448 | 0.0452 | 0.0915 | 0.873 | 0.0891 |
| 2.42E-08 | 0.466 | 0.0470 | 0.128 | 0.995 | 0.102 |
| 4.45E-08 | 0.462 | 0.0466 | 0.179 | 1.10 | 0.113 |
| 8.18E-08 | 0.748 | 0.0762 | 0.251 | 1.26 | 0.130 |
| 1.51E-07 | 0.664 | 0.0675 | 0.351 | 1.44 | 0.147 |
| 2.77E-07 | 0.476 | 0.0481 | 0.492 | 1.68 | 0.173 |
| 5.10E-07 | 0.475 | 0.0480 | 0.689 | 1.86 | 0.191 |
| 9.38E-07 | 0.478 | 0.0483 | 0.964 | 2.38 | 0.245 |
| 1.73E-06 | 0.478 | 0.0483 | 1.35 | 3.23 | 0.333 |
| 3.18E-06 | 0.471 | 0.0475 | 1.89 | 3.87 | 0.399 |
| 5.84E-06 | 0.475 | 0.0479 | 2.65 | 5.15 | 0.532 |
| 1.08E-05 | 0.471 | 0.0476 | 3.70 | 6.17 | 0.637 |
| 1.98E-05 | 0.471 | 0.0475 | 5.19 | 7.15 | 0.739 |
| 3.64E-05 | 0.479 | 0.0484 | 7.26 | 7.59 | 0.784 |
| 6.70E-05 | 0.480 | 0.0485 | 10.2 | 8.58 | 0.886 |
| 1.23E-04 | 0.474 | 0.0478 | 20.3 | 8.95 | 0.925 |
| 2.27E-04 | 0.488 | 0.0493 | 40.7 | 9.43 | 0.975 |
| 4.17E-04 | 0.510 | 0.0516 | 81.3 | 9.57 | 0.990 |
| 7.68E-04 | 0.529 | 0.0536 |  |  |  |

#### 1.2. Experimental single-cell data

The single-cell data that we used were originally presented in ref. (*3*). They are also publically available as example data for the RCell software package, described in ref. (*4*), where this data set is called the ACL394 data set. They are described in the Rcell metadata file (we loaded the data into Rcell and then entered “help(X)” to view the data description) as follows (with typos corrected):

> “This dataset was generated by Cell-ID, from an experiment done in 2004 by Alejandro Colman-Lerner and Andrew Gordon at the Molecular Science Institute (MSI). *Saccharomices cerevisiae* yeast cells of strain TCY3154 (MATa, bar1, prm1::P_Prm1_-YFP::HIS+, trp1::Pact1-CFP::TRP1) were stimulated with different doses of alpha-factor pheromone 10 minutes before the first time point. Images where acquired every 15 minutes for 3.5 hours. In the dataset there are 3 positions per treatment. The strain TCY3154 was derived from ACL394, a W303 derivative.”

The TCY3154 haploid yeast strain was derived from ACLY394, which was derived from ACLY387, which was derived from ACLY379, which is a W303 derivative (the ACLY and ACL prefixes were used interchangeably, so this ACLY379 parent strain is the same as the ACL379 parent of RY2073 described above). The genotype of TCY3154, from the supplementary information of ref. (*3*) is:

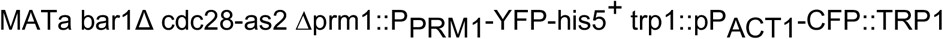

In more detail, TCY3154 is mating type **a**, has the *BAR1* gene removed, expresses Cdc28-as2 from the *CDC28* gene (this replaced the wildtype gene and allows 1-NM-PP1 inhibitor to be added to suppress the activity of the Cdc28 protein, thus stopping progression through the cell cycle), has the *PRM1* gene replaced by a *YFP-His3MX6* gene inserted into the *PRM1* site and under control of the P_Prm1_ promoter, and has a *CFP* gene that is under the control of the constitutive PACT1 promoter and placed in the *trp1* genomic locus.

From ref. (*3*), exponentially growing TCY3154 cells were sonicated, to separate clumped cells, and then deposited into glass-bottomed 96-well sample plates that had been pre-coated with concanavalin-A. Cells were allowed to settle and bind for 10 minutes, after which unbound cells were washed off. Cells were imaged using an inverted fluorescence microscope from which three or more image fields per well were manually selected. Next, the authors acquired time 0 images and changed to medium with α-factor (final concentration of 1.25, 2.5, 5, 10, or 20 nM). Images, at each of the image fields, were collected automatically 10 minutes after that and then every 15 minutes subsequently for a total of 14 time points. The first images in the data set that we used were the ones collected 10 minutes after pheromone addition.

### 2. Hill function fit to experimental dose-response data

#### 2.1. Hill function fitting procedure

Curve fitting to the experimental dose-response data points is described in the supplementary information for ref. (*2*) and repeated here. We fit the YFP to RFP ratio data, listed in the column labeled <y>/<r> in Table SM-1.1, with a 4-parameter Hill function by manually adjusting parameters in Microsoft Excel until the least-squares difference between the curve and the data was minimized. This function is

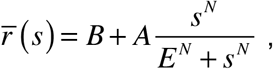

where 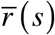 is the mean response over all cells as a function of the signal, *s* is the signal or pheromone concentration, *B* is the Hill function baseline, *A* is the Hill function amplitude, *E* is the Hill function EC_50_ (the signal value that produces half-maximal response), and *N* is the Hill coefficient. We scaled the result so that the baseline would equal the response that arose with no pheromone addition, which was 4.7% of the maximal response, and also so that the fitted response would asymptotically approach a maximum value of 100% (*i.e. A*+*B* = 1.0). Best fit values are *B* = 0.047, *A* = 0.953, *E* = 2.67 nM, and *N* = 1.24. This fit has an rms error of 0.18%, showing excellent agreement with the data.

#### 2.2. Justification for basing results on fits to data rather than the raw data

The channel capacity results found in this work were based on multiple fits to data rather than the raw data themselves. The most important reason for this approach was that it produced an SRV curve that was continuous over all signal values rather than being defined at only the 5 pheromone concentrations that were experimentally investigated for the single-cell data. Using the raw data would have artificially lowered channel capacity results. In particular, it would have capped channel capacity results to log_2_(5) = 2.3 bits (the signal entropy), even if the variation had been very low. The interpolations and extrapolations that were inherent to the fits removed these artifacts. Also, we wanted to quantify the channel capacity during steady-state signaling, which is accurately represented in the dose-response curve that we used but would be complicated to infer from the single-cell data due to time-dependent dose-response behaviors in the PRS (*5*).

### 3. Initial processing and filtering of experimental single-cell data

#### 3.1. Removing dead cells and segmentation errors

Microscope images for the experimental single-cell data were processed in several steps. First, Colman-Lerner, Gordon, and coworkers imported them into CELL-ID for image segmentation and fluorescence quantification (*3, 6*). The CELL-ID output was called the ACL394 data set and was made available with the Rcell download package (*4*).

We imported the ACL394 data set into Rcell, where we applied minimal filtering while following the procedure recommended in ref. (*4*). This removed “badly found and spurious cells,” which are image structures that CELL-ID incorrectly identified as cells, such as dead cells and portions of cells that CELL-ID segmented incorrectly. The first filter removed cells that had highly non-circular images. The following figure shows the distribution of non-circularity values (called fft.stat) for cell images in the ACL394 data set (the figure is copied from the Rcell.pdf document, from the Rcell download package). It shows that nearly all cells had non-circularity values of 0.5 or less, while a few were more non-circular. Rcell removed images with non-circularity values greater than 0.5, which was 1.1% of the images.

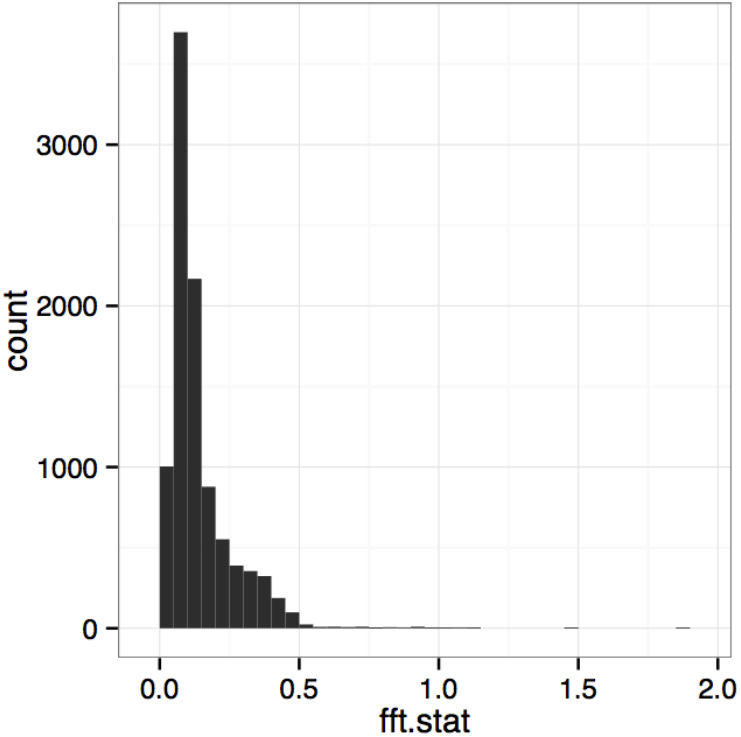

The second filter removed cell images that appeared in only some but not all of the 14 time frames, again because this is indicative of badly found and spurious cells (*4*). This filter removed 21.5% of the cell images. See the Rcell.pdf file, from the Rcell download package, for details. The resulting filtered data are called the ACL394filtered data set and are also available as part of the Rcell package.

#### 3.2. Correcting fluorescence values

Using the ACL394filtered data set, we subtracted off the background fluorescence values for both the YFP and CFP images by following the advice given in the transform.pdf file, from the Rcell download package. In particular, we entered “X<-transform(X,f.total.y=f.tot.y-f.bg.y*a.tot)” in R for the YFP data, where this means that we created the variable f.total.y and set it equal to the raw fluorescence value, f.tot.y, minus the product of the background fluorescence level, f.bg.y, and the cell area, a.tot. Similarly, we corrected the CFP data with the statement “X<-transform(X,f.total.c=f.tot.c-f.bg.c*a.tot)”. We exported the resulting cell fluorescence values from R and imported them into Mathematica.

At this point, our data represented the experimental YFP and CFP fluorescence values, integrated over entire cell areas and corrected for background fluorescence, for each of about 100 cells at 14 different time points, for 5 different pheromone concentrations. The following table (Table SM-2.1) shows the actual number of cells in the ACL394filtered data set at each pheromone concentration (the same cells were investigated repeatedly for the 14 time points, but different cells were used for the 5 pheromone concentrations).

**Table SM-1.2.** Filtering to remove badly found and spurious cells.

| α-factor<br>(nM) | number of cells in<br>ACL394filtered |
| --- | --- |
| 1.25 | 100 |
| 2.5 | 114 |
| 5 | 113 |
| 10 | 117 |
| 20 | 103 |

#### 3.3. Removing cells with outlier data points

These data still contained cell fluorescence values that were clearly in error. For example, the following figure shows YFP fluorescence values over time for the first 20 cells in the data set that were exposed to 20 nM α-factor, with a separate trace for each cell.

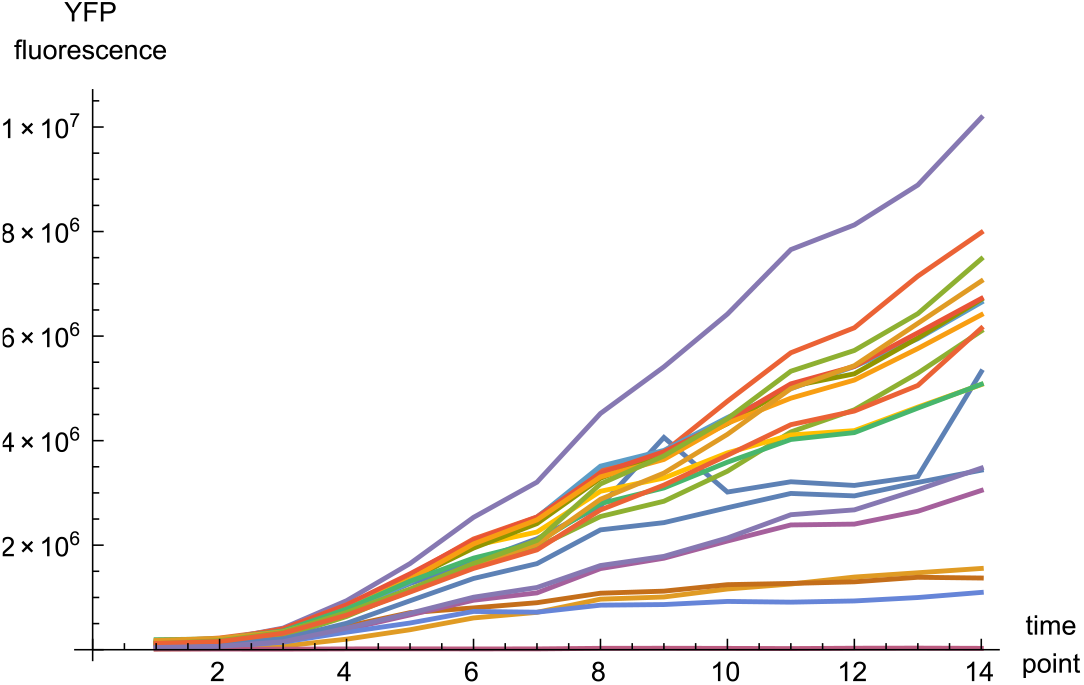

Clearly, most of these cells show a reasonably smooth increase of YFP fluorescence, as one would expect (YFP is expressed by pheromone-responsive genes and is not degraded to any significant extent). However, the cell that is shown in blue near the center of the group of lines appears to have YFP fluorescence that increased suddenly between time points 8 and 9, and then decreased a similar amount between points 9 and 10, and then increased again between time points 13 and 14. This does not make physical sense, but presumably arose from imaging errors at time points 9 and 13.

We removed cells that exhibited this type of artifact from the data set. For each cell, we computed the change in fluorescence between each pair of adjacent time points and then subtracted the mean slope of the time course; we labeled these values as ΔYFP and ΔCFP fluorescence. For example, the same 20 single-cell time traces shown above led to the following figure for the ΔYFP fluorescence values:

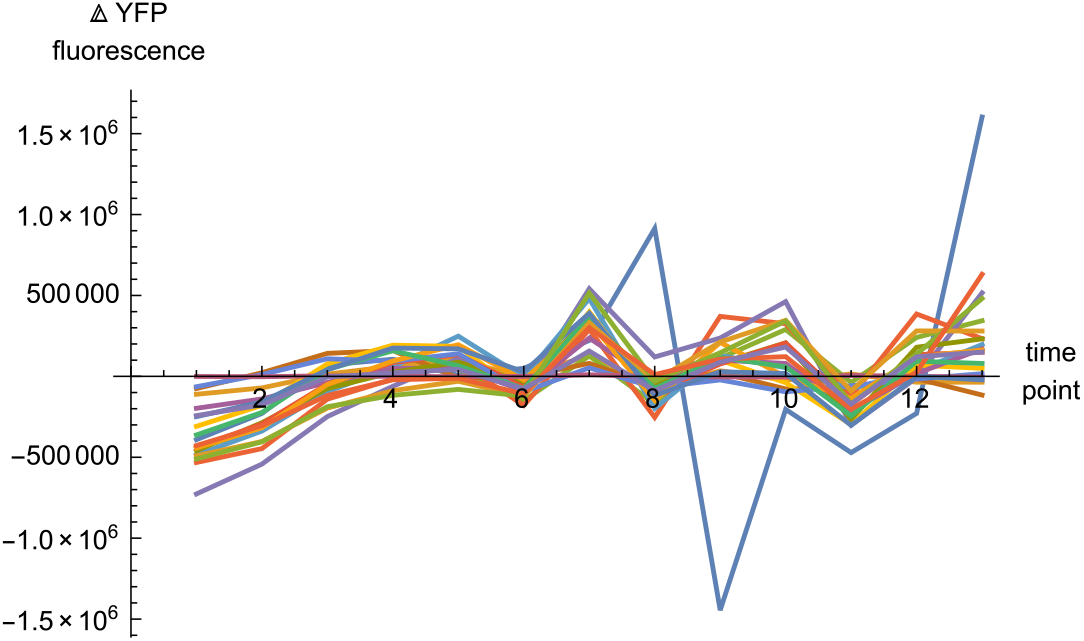

Again, the blue trace stands out as an outlier. We addressed it, and others like it, by removing all cells from the data set for which the absolute value of any ΔYFP or ΔCFP value was in the top 1% of all difference values. For example, the following histogram shows the distribution of difference values for the YFP data at 20 nM α-factor. 1% of these data were outside of the range from −1.003×10^6^ to 1.003×10^6^ fluorescence units, so we removed the cells that contributed those data points from the data set. Looking back at the previous figure, it can be seen that two points of the blue trace shown above had ΔYFP values that exceeded this threshold and no points for the other traces exceeded it, in agreement with the qualitative assessment that the blue trace is an outlier and the others are not.

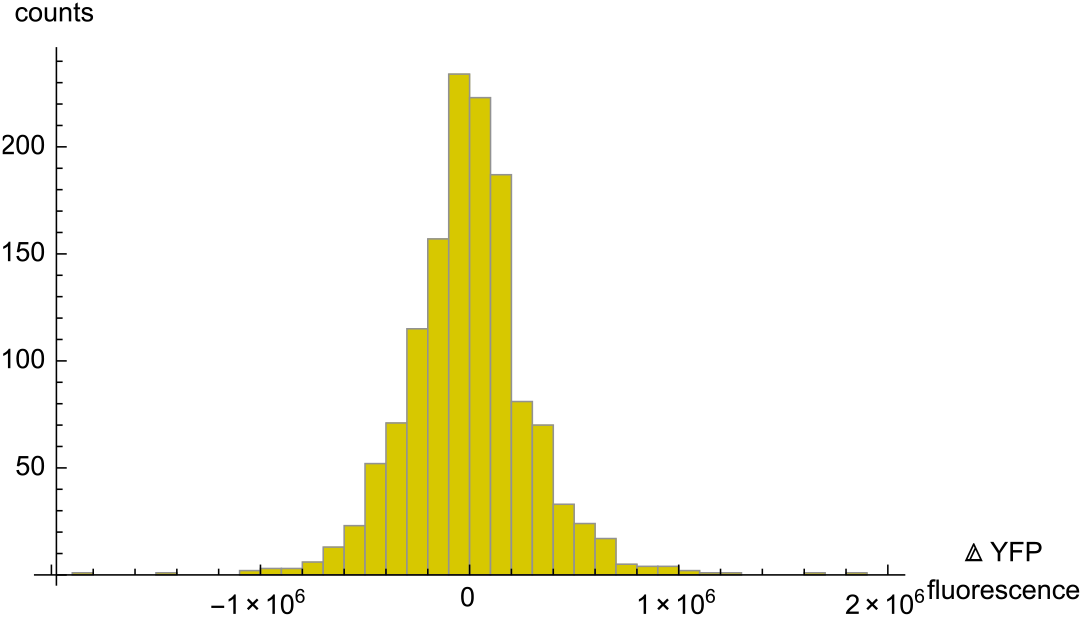

The particular thresholds were different for the other pheromone concentrations and for the CFP data, but in all cases they separated the overwhelming bulk of the distribution from extreme outliers. This filtering removed about 11% of the cells from the data set. The following table shows the numbers of remaining cells at each pheromone concentration.

**Table SM-2.2.** Filtering to remove extreme outlier cells.

| α-factor<br>(nM) | number of cells in<br>ACL394filtered | number of cells after<br>outlier filtering |
| --- | --- | --- |
| 1.25 | 100 | 87 |
| 2.5 | 114 | 105 |
| 5 | 113 | 102 |
| 10 | 117 | 104 |
| 20 | 103 | 90 |

### 4. Population SRV curve computation

#### 4.1. Un-normalized data and coefficients of variation

The following figures show the filtered YFP (left) and CFP (right) single-cell fluorescence data at 1.25, 2.5, 5, 10, and 20 nM α-factor, respectively. The *x*-axis shows the time in minutes and the *y*-axis shows fluorescence in arbitrary fluorescence units.

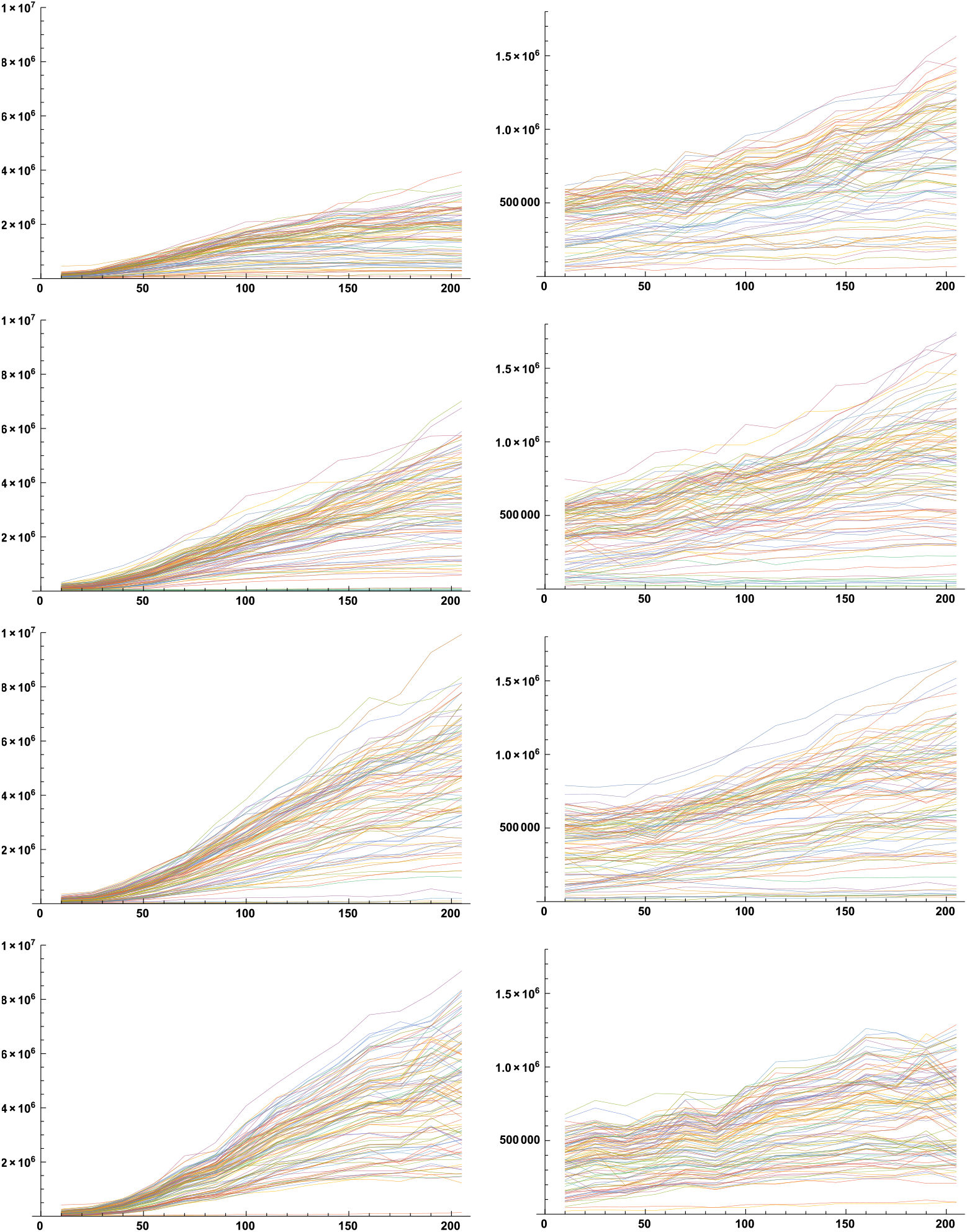
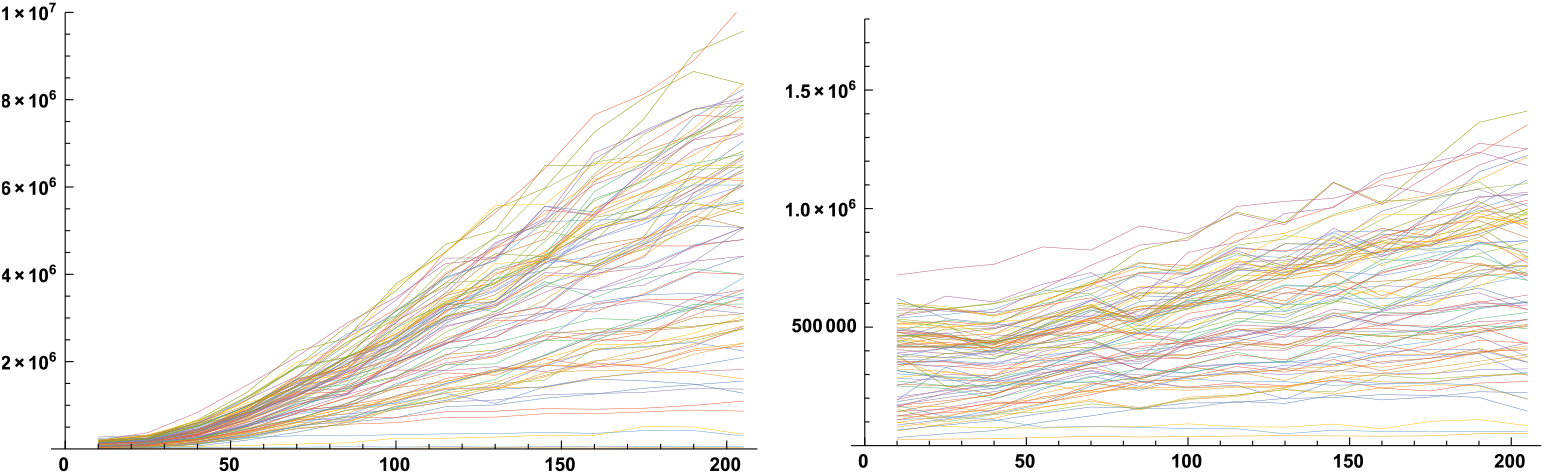

As described in the main text, we computed the mean and standard deviation fluorescence values for each time point and each pheromone concentration, in each case averaging over all cells measured for that time point and pheromone concentration. This led to 70 (5×14) data points for the YFP data, which we call the population mean and standard deviation values. Defining *y*(*s*,*i*,*t*) as the YFP fluorescence value for pheromone level *s*, cell number *i*, and time point number *t*, the population mean and population standard deviations are, respectively

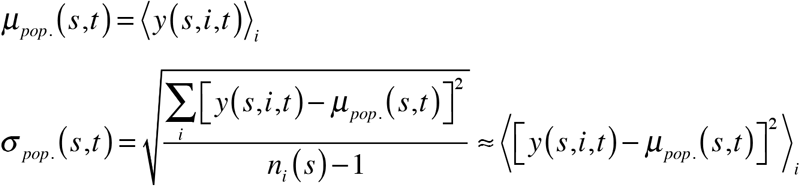

where angle brackets denote a mean over the variable that is listed in the subscript and *ni*(*s*) is the number of cells that were investigated for pheromone level *s* (roughly equal to 100).

Although not described in the main text, we also did the same thing with the CFP data, producing 70 more data points. The following figure shows both the YFP and CFP data, with YFP in yellow and CFP in cyan.

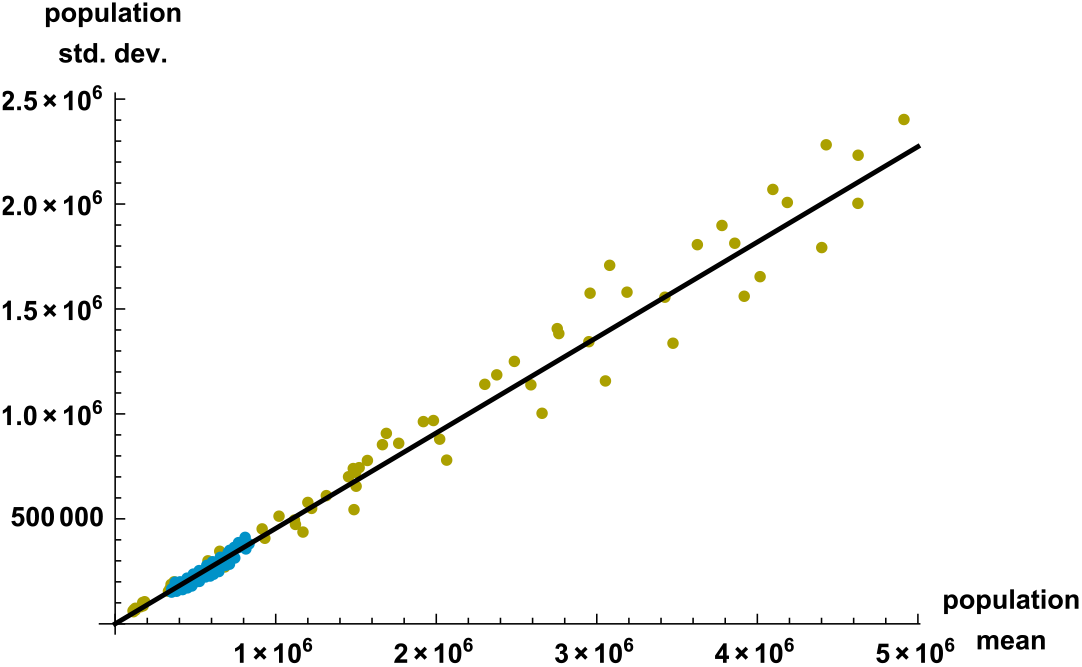

This figure shows that the YFP data exhibited a wider variation of both mean and standard deviations than the CFP data, but that they had almost identical coefficients of variation. Best-fit lines, which were constrained to go through the origin, had slopes (*CV_pop_.* values or, equivalently, η_pop_.) of 0.466 for YFP and 0.443 for CFP. We also fit these lines with unconstrained *y*-intercepts and found that the best-fit intercept parameters were not statistically significant (*p*-values of 0.75 for YFP and 0.22 for CFP), so we neglected these intercepts in further work. Although the similarity between YFP and CFP was interesting, we did not make use of it in this work, but focused primarily on the YFP data. We investigated the scatter of YFP and CFP points about the best-fit lines in this figure and did not find significant trends with respect to either the time after pheromone addition or pheromone dose.

The *CV_pop_.* values of 0.466 and 0.443 are comparable to those found in prior work, which generally ranged from about 0.2 to about 0.7 (*3, 7, 8*).

#### 4.2. Normalized data and bimodal distribution of normalized fluorescence

The following figures show the normalized YFP (left) and CFP (right) single-cell fluorescence data at 1.25, 2.5, 5, 10, and 20 nM α-factor, respectively. Data were normalized by subtracting the mean fluorescence value from each data point and then dividing by the standard deviation, where both means and standard deviations were computed separately at each time point and pheromone dose. As a result, the normalized data have a mean of zero and a standard deviation of 1 at each time point and pheromone dose, when considering the statistics over all cells. As equations, the normalized data values are

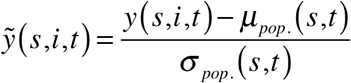

where the tilde denotes normalization. In these figures, the *x*-axis shows the time in minutes and the *y*-axis shows normalized fluorescence.

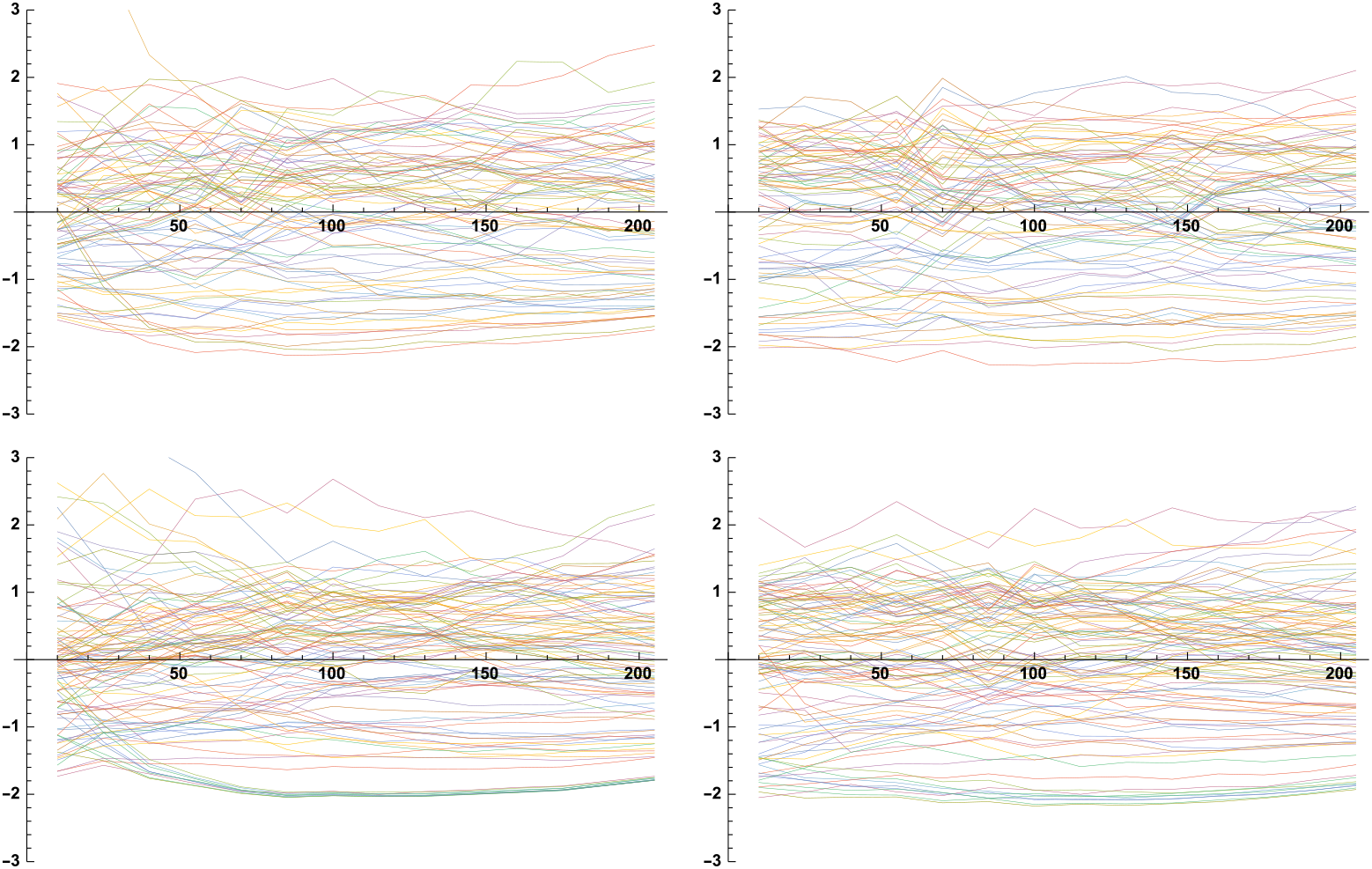
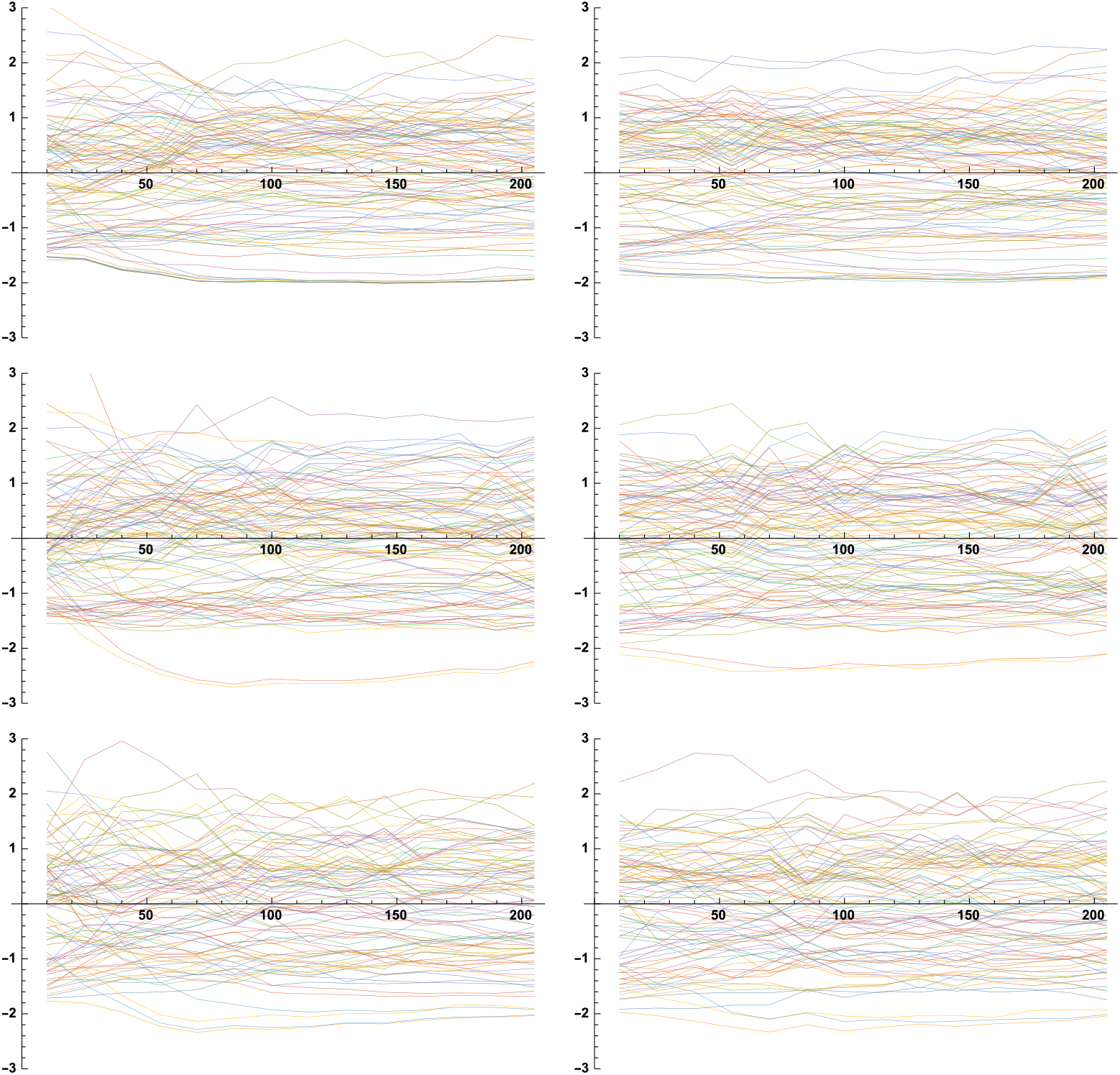

We computed the distribution of these normalized points about the population mean values (equal to 0 in the normalized data) from these data. As described in the main text, the distribution for all of the normalized YFP data points was strongly bimodal and fit well by a sum of two Gaussians. The best-fit parameters for these Gaussians were: area of 0.708, mean of 0.458, and standard deviation of 0.656 for one, and area of 0.283, mean of −1.200, and standard deviation of 0.488 for the other. Further investigation showed that this distribution varied some with different pheromone concentrations and measurement time, but was always bimodal and reasonably similar. For example, the following figures show the distributions at 1.25, 5, and 20 nM α-factor, in each case including values from all time points. Each figure shows the same sum of Gaussians that best fit the complete data set.

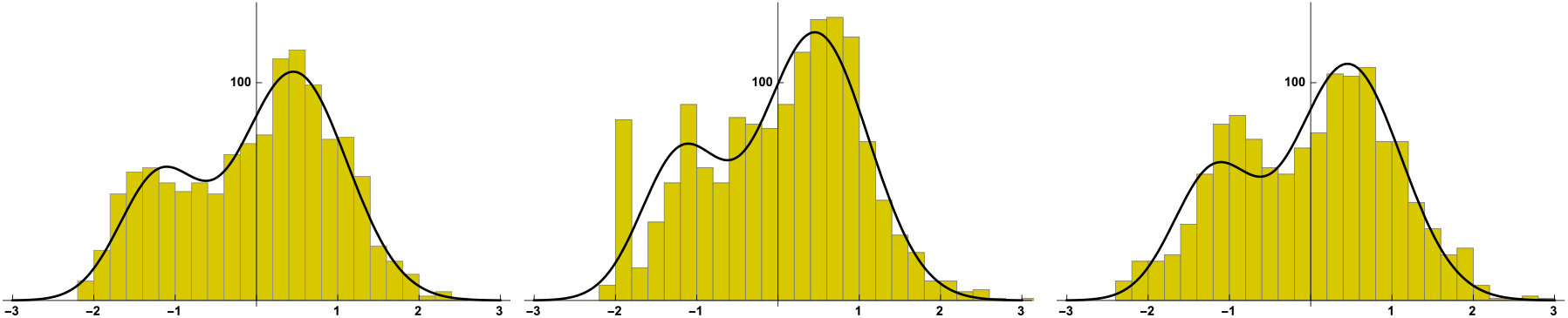

Similarly, the following figures show the distributions at 10 minutes, 115 minutes, and 220 minutes (time points 1, 7, and 14) after pheromone addition, in each case including values from all pheromone doses. Again, each figure shows the same sum of Gaussians that best fit the complete data set.

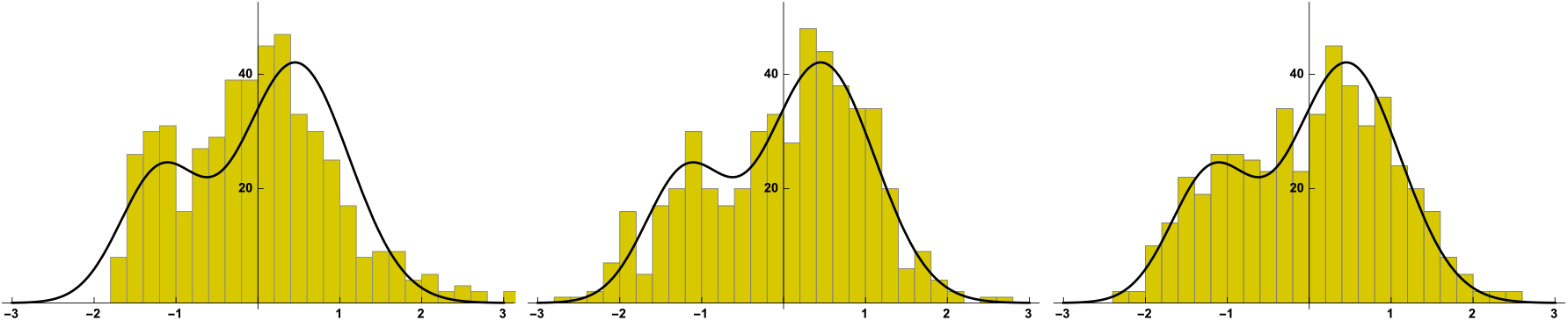

In addition, the distribution for normalized CFP fluorescence values was bimodal as well. It is shown in the following figure, now including all time points and pheromone doses, again with the black line showing the same sum of Gaussians.

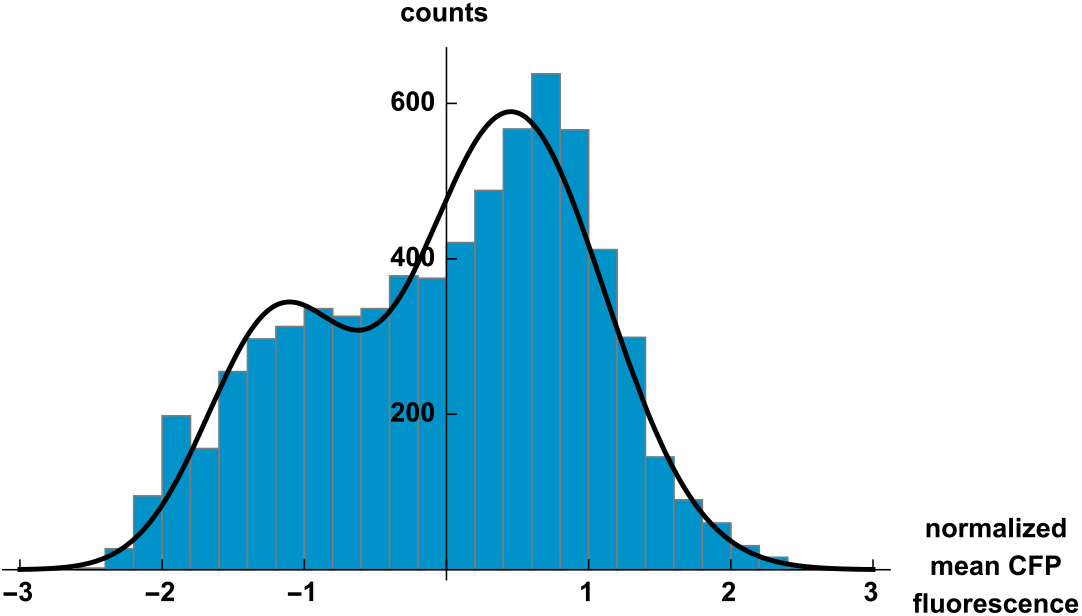

We suspect that this bimodal distribution arose from cells that were arrested in the G1 and G2 phases of the cell cycle.

#### 4.3. Signal-response variation curve

We computed the signal-response-variation (SRV) curve as described in the main text, using the Hill function fit to the steady-state dose-response data (SM-2), the measured coefficient of variation for YFP (SM-4.1), and the fit to the bimodal YFP fluorescence distribution (SM-4.2). We repeat the SRV curve here:

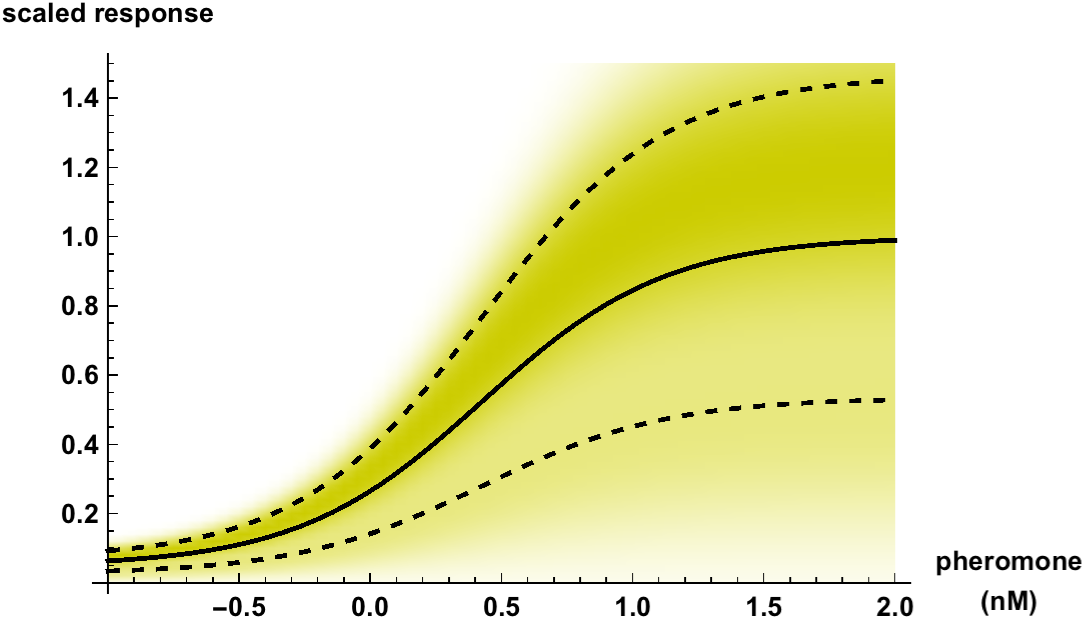

Shading in this figure qualitatively represents the conditional probability that some specific cell will respond with response (fluorescence) *r*, given that the signal (pheromone) level is *s*, written as *p*(*r*|*s*). However, the shading was scaled in this figure to give the same maximum darkness at each signal level. In fact, the conditional probability integrates to 1 at each signal level,

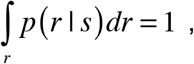

so it would be more correct to show this figure with the same cumulative darkness at each signal level. However, doing so would lead to extremely dark shading at low pheromone concentrations and imperceptibly faint shading at high pheromone concentrations, making the figure difficult to interpret. Thus, we chose this alternate scaling method for clarity.

### 5. Channel capacity computation

#### 5.1. Conditional probability function, p(r|s)

We computed the channel capacity from the *p*(*r*|*s*) function, which is the conditional probability of observing response *r*, given that the signal value is equal to *s*. The signal-response-variation (SRV) curves that we presented are graphical depictions of this conditional probability distribution. We defined the *p*(*r*|*s*) function from three components.

First, 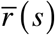 is the mean response for given signal, averaged over many cells, meaning that it is the conventional dose-response curve. In practice, we used

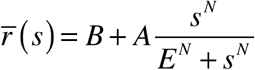

where *B* = 0.047, *A* = 0.953, *E* = 2.67 nM, and *N* = 1.24, as described in the main text and explained further in SM-2. The second component is the coefficient of variation, written here as η (it is denoted *CV* in the main text, but is clearer in equations as *η*). In principle, the coefficient of variation could have been a function of the mean response level and/or the signal value, but the experimental results showed that it was actually a constant (SM-4.1), so it is shown that way here. Third, we defined *f*(*r’*,σ) as the distribution of individual responses about the mean response, scaled so that its standard deviation is equal to σ. As equations, this means that the moments of the distribution of *f*(*r’*,σ) are

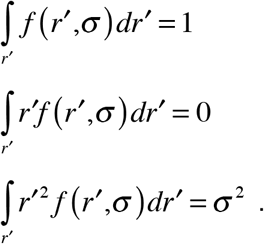

In practice, we used the following bimodal distribution when computing the conditional probability for the cell population,

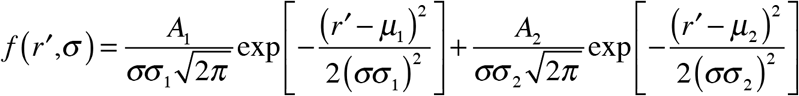

where *A*_1_ = 0.708, μ_1_ = 0.458, σ_1_ = 0.656, *A*_2_ = 0.283, μ_2_ = −1.200, and σ_2_ = 0.488. This bimodal distribution is identical to the one shown in the main text Figure 2D and described in SM-4.2 when σ = 1, and is re-proportioned to be narrower or wider with smaller or larger σ values. When computing the conditional probability for single cells, we used a standard normalized Gaussian distribution,

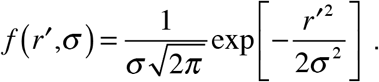

Both of these functions obey the moments that are specified above. Finally, we combined these components to give the conditional probability distribution as

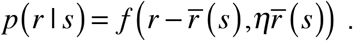

#### 5.2. Principle of channel capacity computation

The mutual information between signal and response cannot be computed from the conditional probability distribution alone but also requires the signal distribution, *p*(*s*). Ideally, the distribution of signals that yeast actually observe in nature would be known, and could be used for *p*(*s*). However, this natural signal distribution is not known, so an alternative approach is to find the *p*(*s*) function that maximizes the mutual information. At this optimum, the mutual information is called the channel capacity. Because of the difference between natural signal distributions and the optimal ones, channel capacity values are upper limits to the mutual information that is actually transmitted by a natural signal. There are also some benefits to focusing on the channel capacity rather than the mutual information that occurs in nature. In particular, the fact that it doesn’t make any assumptions about the natural signal distribution removes a potential source of error and also makes it easier to compare different results with each other.

Vice versa, calculated channel capacity results are also lower limits for the true channel capacities because signaling noise that is measured includes contributions from the actual signaling noise within a cell and experimental noise.

To optimize the mutual information over the possible signal distribution functions, one needs to start with an initial signal distribution that can be improved upon. We defined this initial signal distribution, denoted *p*^1^(*s*), as

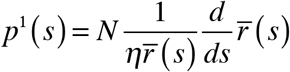

where *N* is a normalization constant set so that

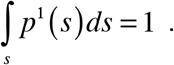

The first term of this signal distribution definition makes the probability inversely proportional to the amount of noise, so signals that are subject to lower noise are used preferentially. The second term of the initial signal distribution equation makes the probability proportional to the slope of the dose-response curve because this leads to a uniform probability distribution over the responses. This latter procedure, known as histogram equalization in digital image processing, optimizes the information transfer in the case of uniform noise (*9*). This initial signal distributions is equal to the optimal signal distribution, meaning the one that gives the greatest possible mutual information, if, and only if, the variation is much less than the mean response (*10*).

The mutual information between signal and response, *I*(*s*;*r*), is (*11*)

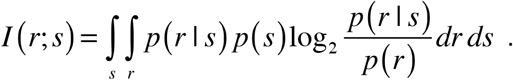

where *p*(*r*) is called the response probability distribution and represents the probability density of observing response *r* while considering all signal values. It is

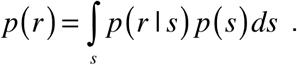

We iteratively optimized the signal distribution using the Blahut-Arimoto algorithm. As explained in ref. (*12*), the Blahut-Arimoto algorithm is based upon a variational optimization of the mutual information. Its procedure involves the alternation of two calculations. First, the signal distribution is updated from iteration *n* to *n*+1 with the equation

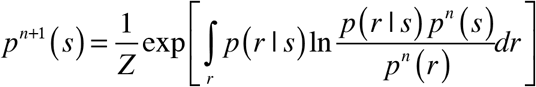

where *Z* is a normalization constant, defined so that

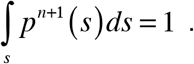

Second, the response distribution is updated using the same equation as above,

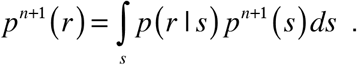

While it is possible to constrain the signal distribution during the optimization, such as to add a cost to particular signals or to enforce a certain degree of smoothness (*12*), we did not do so in this work. As a result, we computed the global optimal channel capacity, rather than values that depended upon essentially arbitrary choices of constraints.

#### 5.3. Numerical channel capacity computation

In practice, these integrals need to be computed numerically. We did so, in the Mathematica software, by partitioning signal values from 10^-4^ to 10^4^ (nM of pheromone) with a logarithmic step size of 0.05 (e.g. possible signal values were at 10^-4^, 10^-3.95^, 10^-3.9^,…, 10^3.95^, 10^4^); these are equally spaced along the logarithmic dose axes that are conventionally used to show dose-response curves. We also partitioned response values from −1 to 4 in linear steps of 0.05. Negative response values are physically nonsensical but are mathematically necessary because the assumption of Gaussian variation makes negative responses possible, although improbable (this is another reason why the channel capacity computations are upper limits for the actual mutual information). We tabulated the *p*(*s*) function using the signal value partitions, the *p*(*r*) function using the response value partitions, and the *p*(*r*|*s*) function as a matrix using both partitions. The following figures show the initial signal distribution, the initial response distribution, and *p*(*r*|*s*), all for the channel capacity computation for the cell population:

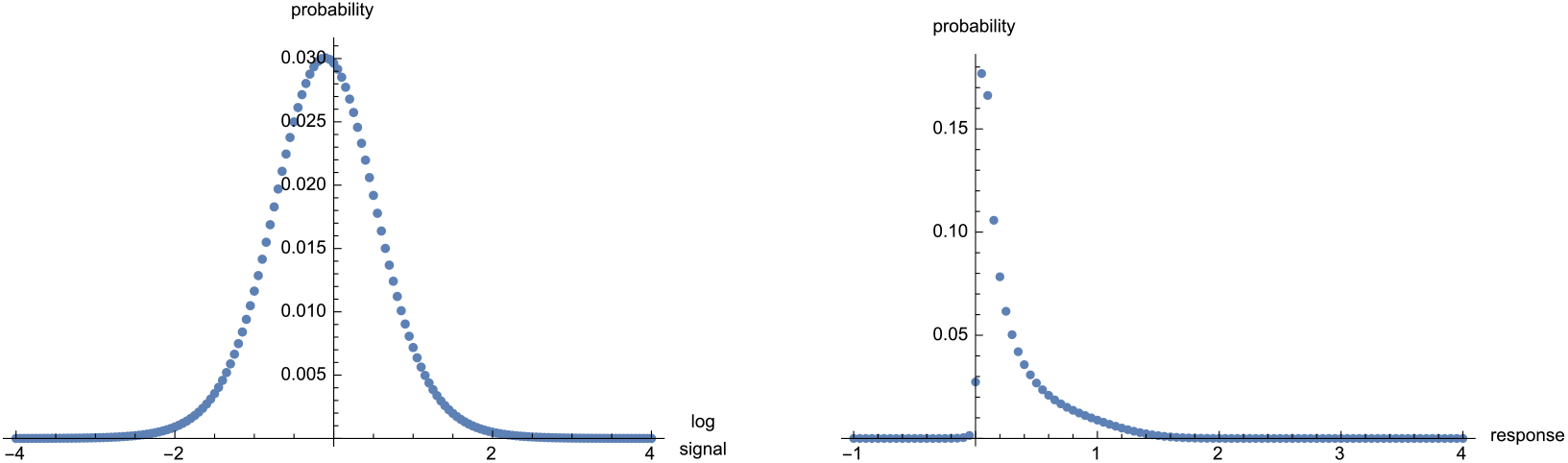

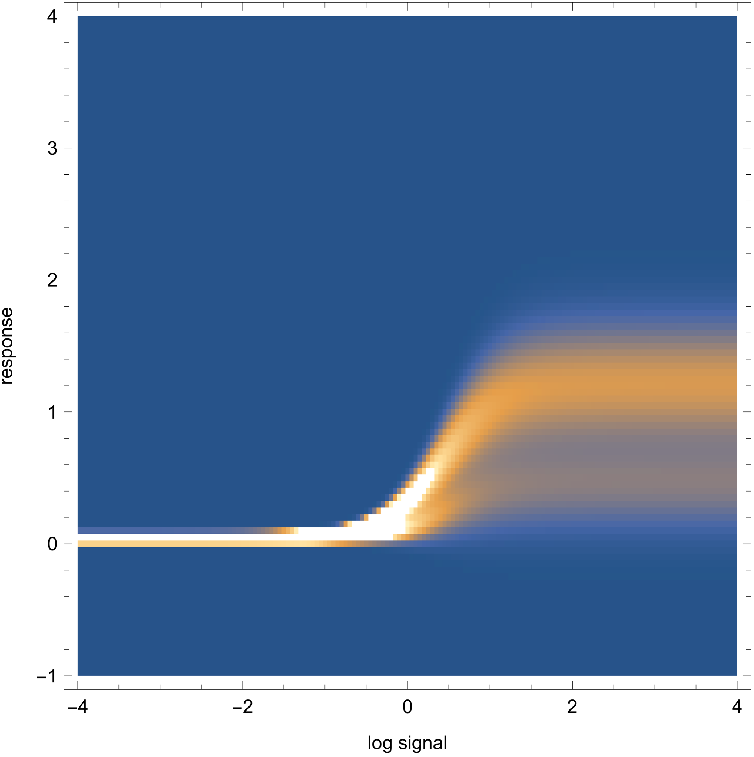

We optimized the signal distribution using discrete versions of the above equations:

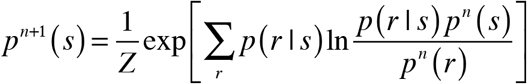

where *Z* is defined so that

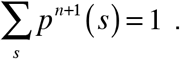

Also,

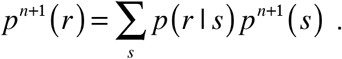

We recorded the mutual information during the optimization, computing it from the equation

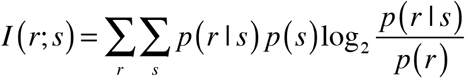

We performed 200 iterations of optimization, at which point the mutual information had nearly stopped changing, shown with blue points in the following figure:

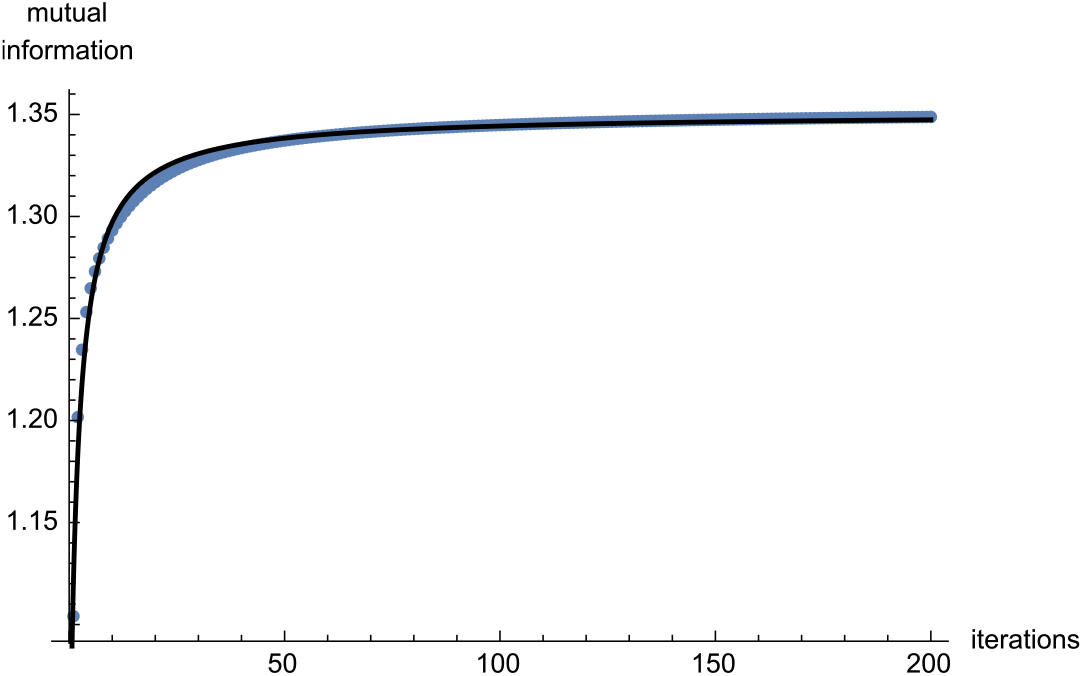

The amount of mutual information increase during optimization depended upon the amount of variation but was always reasonably modest. In particular, it increased from 1.10 to 1.35 bits for the cell population calculation, where the variation was large, and it increased from 2.52 to 2.66 bits for the single-cell calculation, where the variation was much smaller. The mutual information did not quite stop changing even after 200 iterations, so we fit these mutual information data points to a rational function of the form

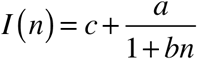

where *n* is the iteration number and *a*, *b*, and *c* are fit parameters. Taking this equation to the limit of infinite iterations shows that *I*(∞) = *c*, meaning that *c* is the channel capacity. These fit parameters are the channel capacity values that we presented in the main text.

Interestingly, the signal distribution evolved substantially during the optimization, changing from a simple smooth unimodal distribution to a widely spaced and spiky distribution. It also did not stop evolving with more iterations, but continued becoming more widely spaced and more spiky, as seen in the following figure,

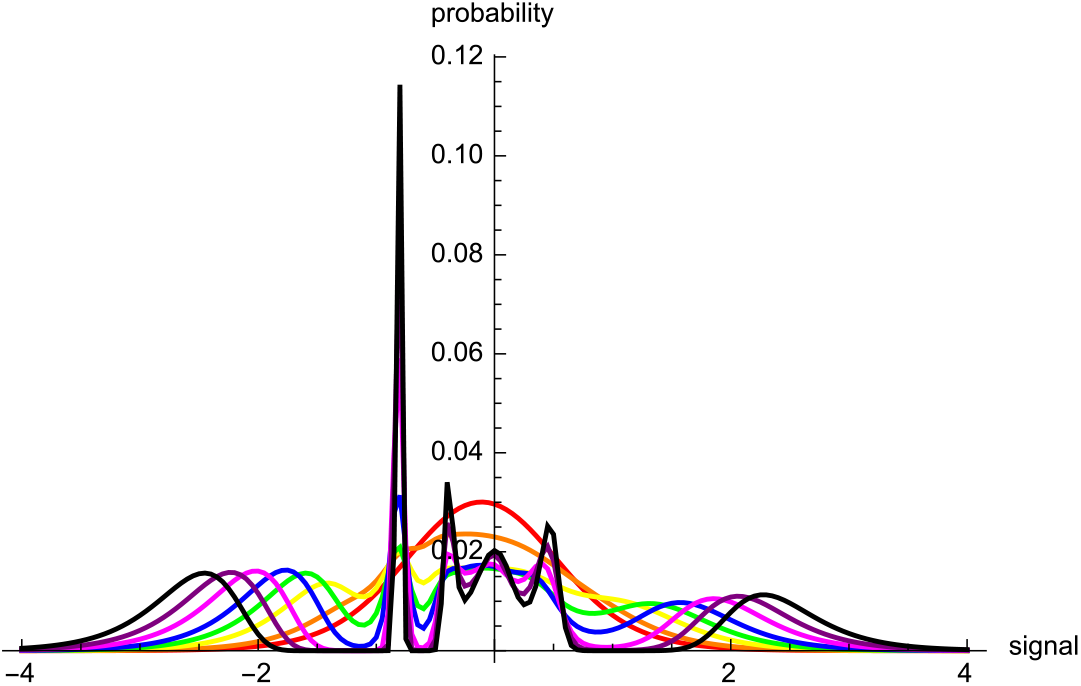

The lines show the signal distribution at iteration 1 (red), 2 (orange), 5 (yellow), 10 (green), 20 (blue), 50 (magenta), 100 (purple), and 200 (black). Note that the channel capacity for this calculation was only 1.35 bits, which corresponds to 2^1.35^ = 2.5 distinguishable signals. However, the final signal distribution has 6 peaks in it and would likely have many more if we had continued the optimization.

In contrast to the signal distribution, the response distribution did not change appreciably after the first several iterations, as shown below,

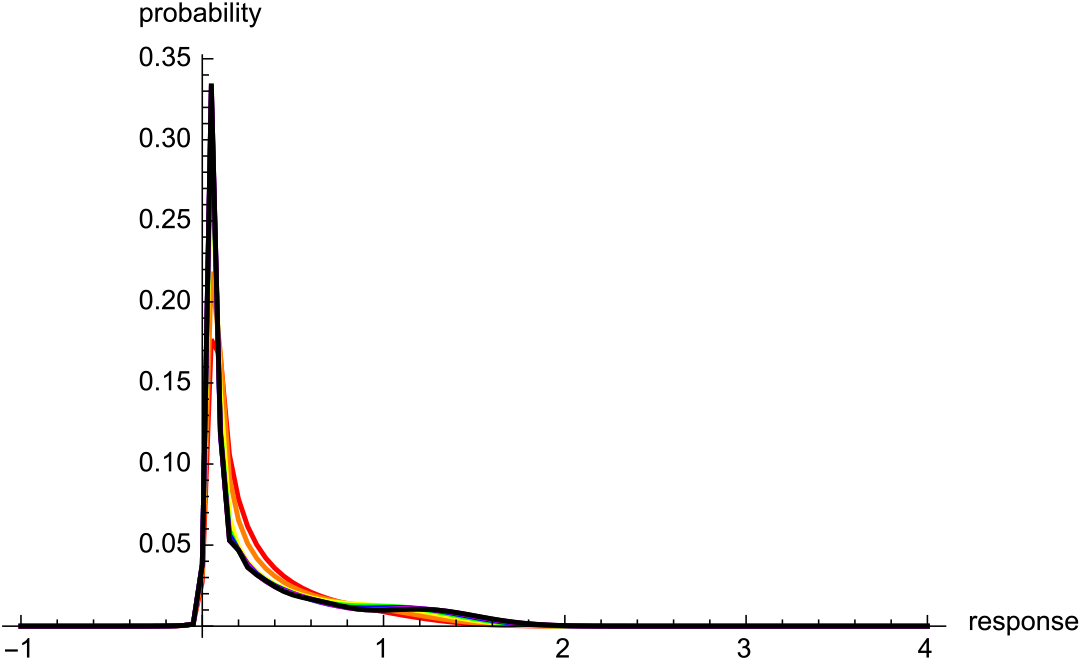

The lines and colors are the same as for the signal distribution. The fact that neither the response distribution nor the mutual information changed substantially during optimization, even as the signal distribution went from a smooth unimodal distribution to a very spiky one, indicates that the precise shape of the signal distribution is not particularly important. Thus, the fact that we did not constrain the signal distribution during optimization cannot have made a substantial difference to the computed channel capacity.

### 6. Single-cell SRV curve computation

We computed the single-cell variation in a few different ways, all of which yielded similar results. The method shown in the main text is arguably the best method, although the others are somewhat simpler and also informative so we describe them here as well. This section presents some mathematical definitions first, then the other two methods, and finally details of the method that is presented in the main text.

#### 6.1. Normalized single-cell noise values

The normalized fluorescence traces (main text Fig. 2C) generally have minimal overall slope but instead show noise about relatively constant values. To compute this noise, we started by quantifying the mean value of each cell’s normalized fluorescence. The normalized single-cell mean value for cell number *i* at pheromone level *s* is

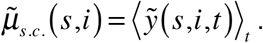

The following figure shows a black line that is a representative set of normalized fluorescence values for a single cell and a solid blue line that represents the normalized single-cell mean for the same cell. Normalized single-cell noise values are differences between normalized fluorescence values and these mean values,

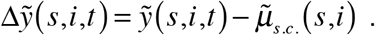

We also calculated the standard deviation of the normalized single-cell fluorescence values about the mean values to yield the normalized single-cell standard deviations.

These values are

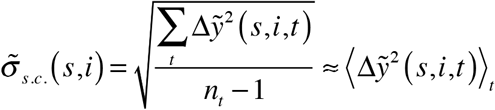

where *n_t_* is the number of time points, and are shown in the following figure with dashed blue lines.

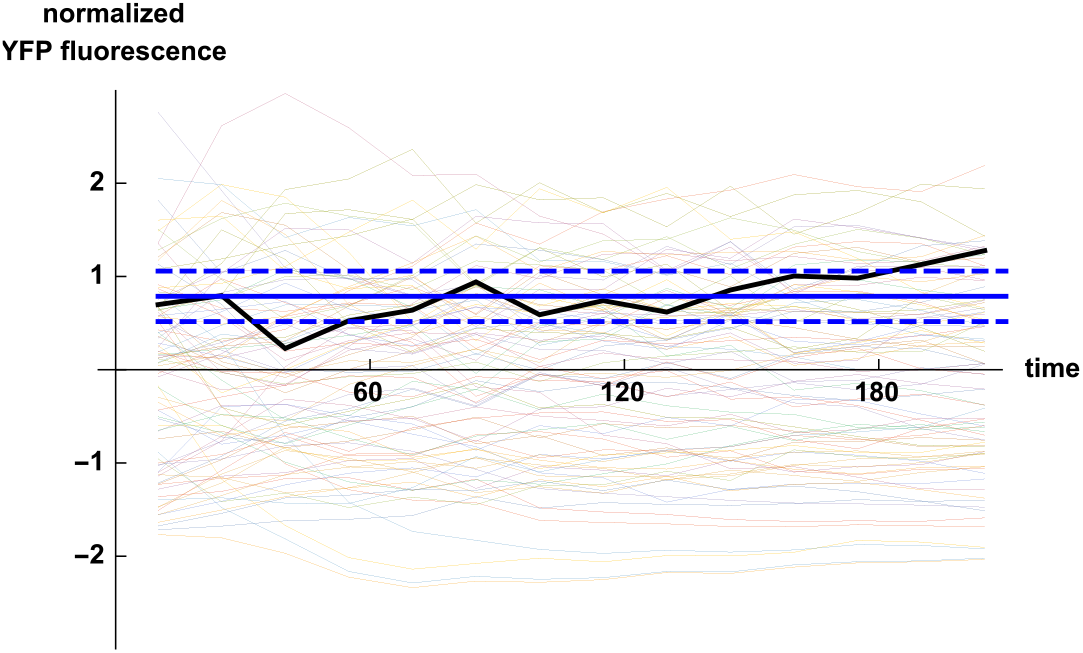

#### 6.2. Mean of normalized single-cell standard deviations

In one approach to quantify single-cell variation, we focused on the normalized single-cell standard deviations, in which the normalized single-cell noise values are averaged over time. The following figures show that the normalized single-cell means and standard deviations exhibited a positive correlation in which the brighter cells exhibited greater fluorescence variability. Each yellow or cyan point represents one cell and results are aggregated for all pheromone concentrations. These figures show that this trend was similar for fluorescence from YFP (left) and CFP (right).

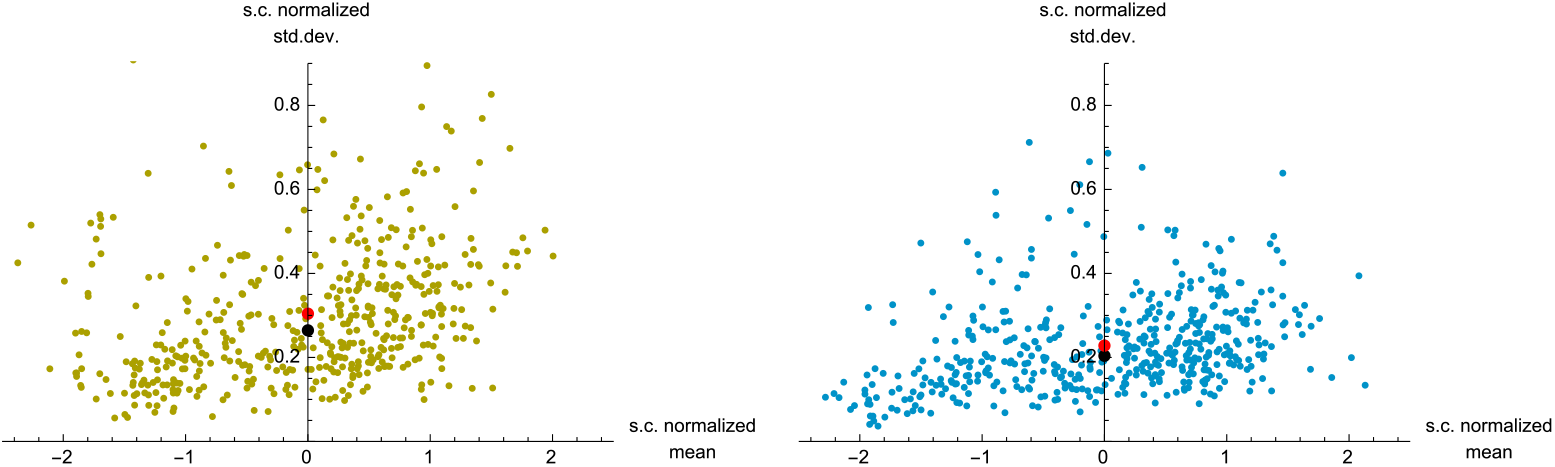

We also graphed these scatter plots at individual pheromone concentrations and found similar distributions of points and similar correlations in all cases.

We defined a “typical” cell as a hypothetical cell that has a normalized single-cell mean fluorescence that is equal to the mean of those values for all cells (equal to 0, due to the normalization process), and that has a normalized single-cell standard deviation that is the mean of those values for all cells. As equations, this typical cell has normalized mean and standard deviation values equal to

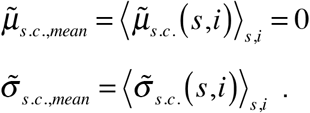

The above scatter plot figures depict this typical cell with red dots. This typical cell has normalized single-cell standard deviation of 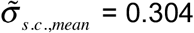 for YFP and, less importantly, 0.228 for CFP. These values are in normalized units, where a value of 1 represents the population standard deviation. A complication is there is a different population standard deviation at each time point and pheromone dose. Thus, we multiplied 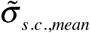 by the population standard deviation, σ_pop._(*s,c*), for each time point and pheromone dose to remove the normalization and convert the value to fluorescence units, yielding a list of single-cell standard deviations, all for the same typical cell. We then divided these standard deviations by the respective population mean fluorescence values (because we assumed that the fluorescence of a typical cell is equal to that of the population mean) to yield the single-cell coefficient of variation for a typical cell. As an equation, these transformations are

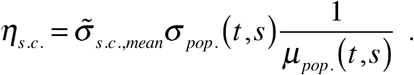

However, we found previously that the population coefficient of variation is independent of time or pheromone dose (SM-4) and so can be written as

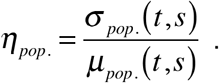

This identity simplifies the single-cell coefficient of variation to

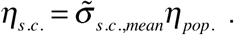

Using this equation, the single-cell coefficients of variation were 0.142 for YFP and 0.101 for CFP. The channel capacity computation described in SM-5 then shows that the YFP coefficient of variation corresponds to a channel capacity of 2.60 bits (The channel capacity cannot be computed for CFP because pheromone does not induce CFP expression).

#### 6.3. Median of normalized single-cell standard deviations

The prior calculation was based on the definition that a typical cell is the one that has the mean amount of fluorescence variation. On the other hand, the median fluorescence variation may be more representative of a typical cell because it treats each of the measured cells equally rather than placing a greater emphasis on ones with unusually large variations; also, some of these large variations likely arose from imaging artifacts that were not caught by the data filtering procedures described above in SM-3.

The median normalized single-cell standard deviations, shown with black dots in the above scatter plot figures, were slightly lower than the mean values due to there being more outliers with high variation than with low variation. In particular, 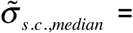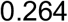 for YFP and 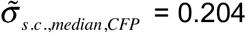 for CFP. This change does not affect the rest of the computation described above, so the single-cell coefficient of variation in this calculation method is

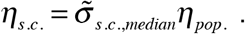

Values were 0.123 for YFP and 0.090 for CFP. The YFP value corresponds to a channel capacity of 2.73 bits, which is slightly larger than the 2.60 bits shown above for a mean cell.

#### 6.4. RMS average noise approach (main text approach)

Both of the prior methods, in which a typical cell is defined as having either the mean or median variation, have the drawbacks that they implicitly assume that the single-cell coefficient of variation is independent of both time and pheromone dose. This occurs because the means and medians are computed from the aggregate of all normalized data values, without considering the possibility of trends over time or pheromone dose. To address this potential problem, we explicitly checked for trends and observed that they were insignificant. Nevertheless, a better approach, and the one that we describe in the main text, is to rearrange the data analysis steps slightly to yield separate single-cell mean and standard deviation values for each time point and pheromone dose. These can then be fit to find the single-cell coefficient of variation, paralleling the approach that we followed when computing the population coefficient of variation.

This procedure started again with the normalized data. Instead of computing standard deviations for individual cells, we instead computed a standard deviation for each time point and pheromone dose, taking one normalized value for each cell and combining these values over all of the cells,

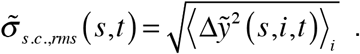

Because this computation combines results from all cells, it still represents the variation for a hypothetical average cell. We then removed the normalization by multiplying by the population standard deviation,

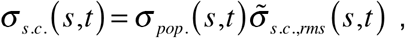

yielding the average noise amounts for each pheromone concentration and at each time point, now in fluorescence units. Meanwhile, we defined the single-cell mean fluorescence values to be the same as the population mean fluorescence values, again because the focus here is on a typical cell,

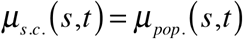

The following figure shows a scatter plot of these single-cell means and standard deviations, with one point for each time value and pheromone concentration. The upper points, in yellow and cyan, are the same ones shown before for the cell population. The lower points, in green for YFP and cyan for CFP, represent the single-cell data.

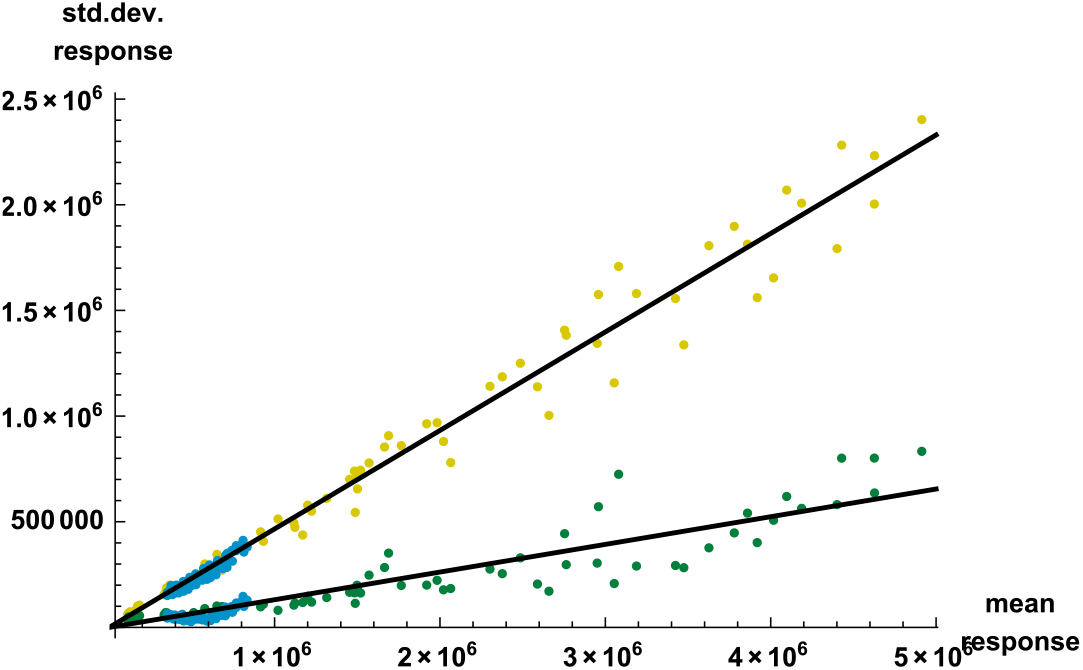

This figure shows a high degree of correlation between single-cell means and single-cell standard deviations, suggesting that the coefficient of variation is a constant value. To confirm this, we fit the data with a line that was not constrained to intersect the origin and found that the *y*-intercept was much less than its standard error and had a P-value of 0.61, both of which showed that the *y*-intercept was insignificant. Fitting these data to line that was constrained to intersect the origin showed that the single-cell coefficient of variation was 0.131 for YFP and 0.104 for CFP. The YFP value corresponds to a channel capacity of 2.66 bits, a value that is very similar to those from the two approaches described above.

### 7. Other sources of variation

#### 7.1. Experimental artifacts

We were unable to differentiate biochemical noise from experimental noise, where this latter source represents the unpredictable errors that arise from the measurement process.

However, we did observe some variation in the data that clearly arose from experimental artifacts. This variation was particularly obvious during the second data filtering process (described in SM-3.3), when we removed cells with outlier fluorescence values. The following figures show the fluorescence difference values (essentially derivatives of the time-dependent fluorescence) for the YFP data on the left and the CFP data on the right for the cells measured at 20 nM α-factor, with one line for each cell. These lines are color coded to represent the three different microscope positions that were used while collecting these data. Clearly, the fluorescence differences are highly correlated for different cells that are at the same microscope position but generally weakly correlated for cells at different microscope positions. Furthermore, the YFP and CFP data show the same trends. For example, the blue traces are all high for both YFP and CFP at time point 9 and all low at time points 8 and 10.

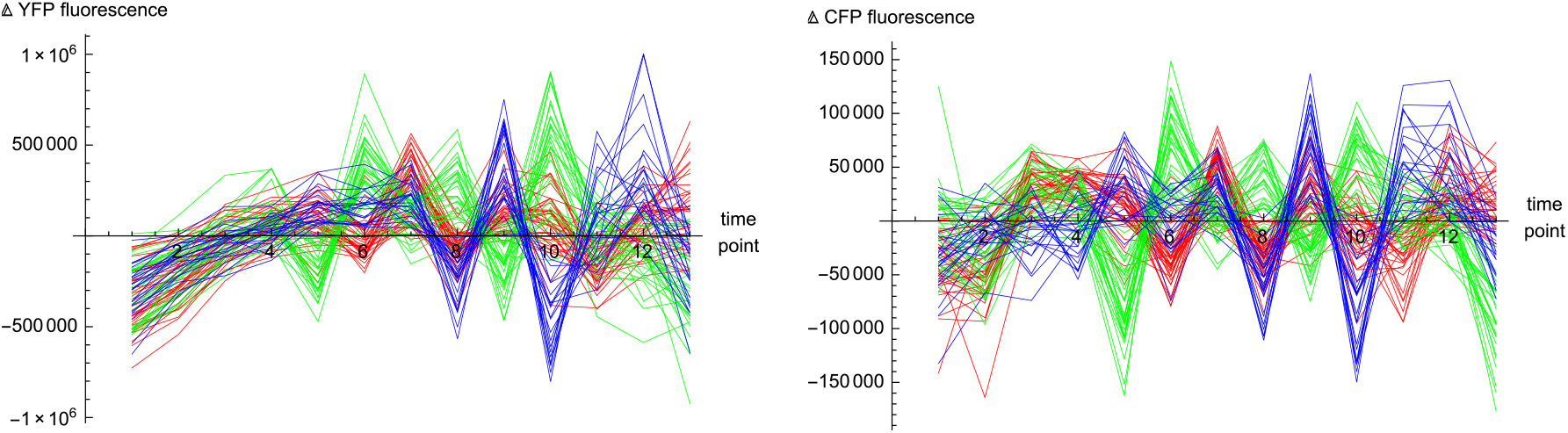

We removed these correlations from the data in preliminary work, but doing so led to additional artifacts and did not affect the results substantially. Thus, we left these artifacts in the data for all of the work described here. Their presence leads to a known source of experimental variation which necessarily decreased the channel capacity calculation results. This implies that the given channel capacity results are upper limits to the true values. These artifacts added variation to the single-cell traces but did not affect cell-to-cell variation significantly, so they undoubtedly had a much larger effect on the single-cell calculation of channel capacity (2.66 bits) than on the population calculation of channel capacity (1.35 bits). On the other hand, they probably did not affect the channel capacity calculation when using internal referencing (2.92 bits) because the CFP and YFP data exhibited similar artifacts and so these would have been removed by the internal referencing procedure.

#### 7.2. Pathway and expression subsystems

The work presented here analyzes data that were collected as part of work presented by Colman-Lerner et al. in ref. (*3*). Those authors analyzed the data in a somewhat different manner, using it to separate the total fluorescence variation observed in yeast cells into its different sources. For cell *i*, they showed that the amount of PRS reporter protein (essentially the YFP fluorescence value, not accounting for experimental noise) at time *ΔT* after pheromone stimulation should equal

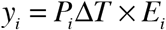

where *P_i_* is the power of the PRS pathway, which is a function of both the pheromone input concentration and the ability of the pathway to transmit signals, and *E_i_* is the power of the cell’s gene expression system. These two power terms can be subdivided into values that are consistent over time for a single cell but might vary between individual cells, which are called capacity terms, and values that vary stochastically over time, which are called noise terms. Thus,

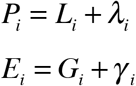

where *L_i_* is the pathway capacity, *λ_i_* is the pathway noise, *G_i_* is the expression capacity, and *γ_i_* is the expression noise. Variation in these values is quantified using the coefficient of variation, η or its squared value, η^2^. The total YFP fluorescence variation for a population of cells, η^2^(*y*), arises from variation in all of these noise terms, so

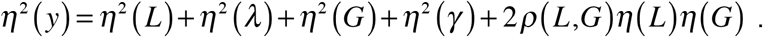

The η^2^ values are added because the variance of the sum of two independent random variables is the sum of their variances. The final term accounts for the possibility that the pathway capacity, *L*, is correlated with the expression capacity, *G*, with correlation coefficient ρ(*L*,*G*); this term is often ignored in practice.

Colman-Lerner et al. used correlations between CFP and YFP fluorescence in two types of experiments, one with both fluorophores induced by pheromone and one with only YFP induced by pheromone, to find that η^2^(*y*) = 0.17, η^2^(*G*) = 0.14, η^2^(*P*) = 0.029, and η^2^(γ) = 0.002 (from their main text p. 702). This means that most of the total observed variation arose from cell-to-cell differences in expression capacity, much less arose from variations in pathway power, and only about 1% arose from expression noise. These values varied some with different yeast strains, over time, and with different pheromone concentrations, but always showed the same trends.

This work quantified two main noise values, the population coefficient of variation, *η*_*pop*_. (SM-4), and the single-cell coefficient of variation, η_*s.c*_. (SM-6), finding *η_pop._* = 0.466 and *η_s.c._* = 0.131. Squaring these values gives η^2^_*pop.*_ = 0.217 and η^2^_*s.c.*_ = 0.017. Taking the ratio of these two numbers shows that 8% of the total variation arises from signaling noise within a cell and 92% of the variation arises from cell-to-cell differences.

All sources of variation contribute to the population variation, so η^2^_*pop.*_ = η^2^(*y*) and, indeed, our result is comparable to the one found by Colman-Lerner et al. The single-cell variation arises from stochastic variation in both the pathway power and the expression system, so η^2^_*s.c.*_ = η^2^(λ) + η^2^(γ). If we accept the result that 1% of the total variation arises from gene expression noise, η^2^(γ), then this implies that 7% of the total variation arises from variation in pathway power, η^2^(λ). On the other hand, Table 1 of Colman-Lerner et al. shows that η^2^(γ) was typically closer to 2% of the total variation, which would then decrease the contribution of η^2^(λ) to about 6%.

Taking these results together shows that total variation can be subdivided into about 82% gene expression capacity, 10% pathway capacity, 6% pathway noise, and 2% gene expression noise.

In part of this work, we considered information transmission when a cell uses an internal standard to calibrate gene expression from a pheromone responsive gene. We investigated this by dividing the YFP fluorescence values by CFP fluorescence values and then computing coefficients of variation and channel capacity values from those results. The population coefficient of variation was *η_pop._* = 0.251 and the single cell coefficient of variation was *η_s.c._* = 0.102, which square to equal η^2^*pop.* = 0.063 and η^2^_*s.c.*_ = 0.0104. Calibration removes the gene expression capacity contribution from this population variation, making it equal to about η^2^_*pop.*_ = η^2^(*L*) + η^2^(λ), where this ignores the portion of η^2^(γ) that is different for different genes within the same cell (intrinsic noise) but keeps the portion that is the same for different genes but varies stochastically over time. Using results found above with just the YFP data, η^2^(*L*) ≤ 0.022 and η^2^(λ) ≤ 0.013, which add to 0.035. The value with calibration, 0.063, is in modest agreement with this prediction but is somewhat larger, probably indicating that the η^2^(*G*) fraction was overestimated slightly. Similarly, calibration makes the single cell variation equal to about η^2^_*s.c.*_ = η^2^(λ), for which the YFP data lead to a prediction of 0.013. In this case, the value with calibration, 0.0102, is similar to the prediction but a little smaller, probably indicating that the pathway noise was overestimated slightly.

### 8. Signaling over time

In general, the cell fluorescence values considered here represent the sum of two components. The first is a deterministic or “true” response that depends on the pheromone dose, the time since pheromone was added, and a cell-specific scaling factor that is constant over time. The second is stochastic noise that arises in the cell signaling pathway, the cell expression machinery, and the experimental measurement. The stochastic noise limits one’s ability to determine the true response, and hence to determine the applied pheromone signal.

If the stochastic noise varies quickly enough that its value at each 15 minute measurement point is independent of its values at prior measurement points, then each measurement would represent an independent observation. In this case, averaging over multiple measurements would improve the estimate of the true response, with the standard error of the estimate decreasing as the square root of the number of measurements. Another way of expressing the same result is that the information that one would learn about the true response would increase linearly with additional measurements.

Real noise values are rarely independent of all prior noise values, but are instead influenced by prior values for some period of time. For example, YFP and CFP have maturation times of about 30 minutes, so high protein expression at one time point leads to increased production of mature fluorescent proteins over, roughly, the next 30 to 60 minutes but has a diminishing influence afterward. In cases where old events are “forgotten” at a constant rate, which is typical, the autocorrelation function of the measured responses decays exponentially, from a value of 1 at a time difference of 0 and approaching 0 for long time differences. The characteristic time of this exponential, τ, is called the autocorrelation time. Noise values become nearly independent of each other every τ time units, so continuous measurements of the response create information about the true response at a rate that is the number of bits for one measurement times 1/τ.

To attempt to quantify τ, we computed the autocorrelation functions for the normalized YFP and CFP data sets, which is shown below.

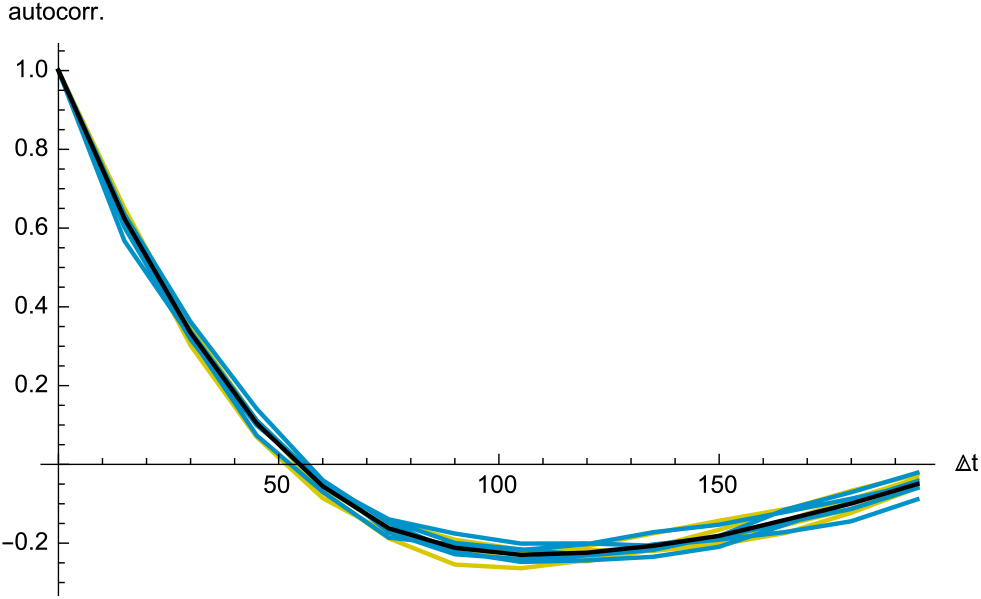

Yellow lines represent autocorrelation functions for YFP data, in each case averaged over all cells and computed separately at each pheromone dose, and cyan lines are similar but for CFP data. The black line represents the average of all YFP and CFP autocorrelation functions.

This curve shape is surprising because it does not show simple exponential decay. Instead, the negative autocorrelation values that have a minimum value at around 2 hours appears to indicate some sort of oscillation, where positive deviations at one time tend to produce negative deviations 2 hours later and vice versa. However, further investigation showed that they are actually an artifact of using a short time series. To both determine this and quantify the autocorrelation time, we created a synthetic data set with 200 “cells”, where each cell’s response at time point *j* was defined as

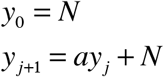

where *N* is a Gaussian distributed random value with zero mean and unit standard deviation and *a* is a “memory” parameter that is equal to zero if prior values are forgotten immediately and 1 if prior values are remembered forever. This function has autocorrelation time τ = −1/ln *a*. We used 14 time points, as for the actual data, and adjusted the *a* parameter until the autocorrelation function for this synthetic data set, shown with the red line in the following figure, matched that of the real data, shown with the black line. Light blue lines represent autocorrelation functions for a few individual model cells.

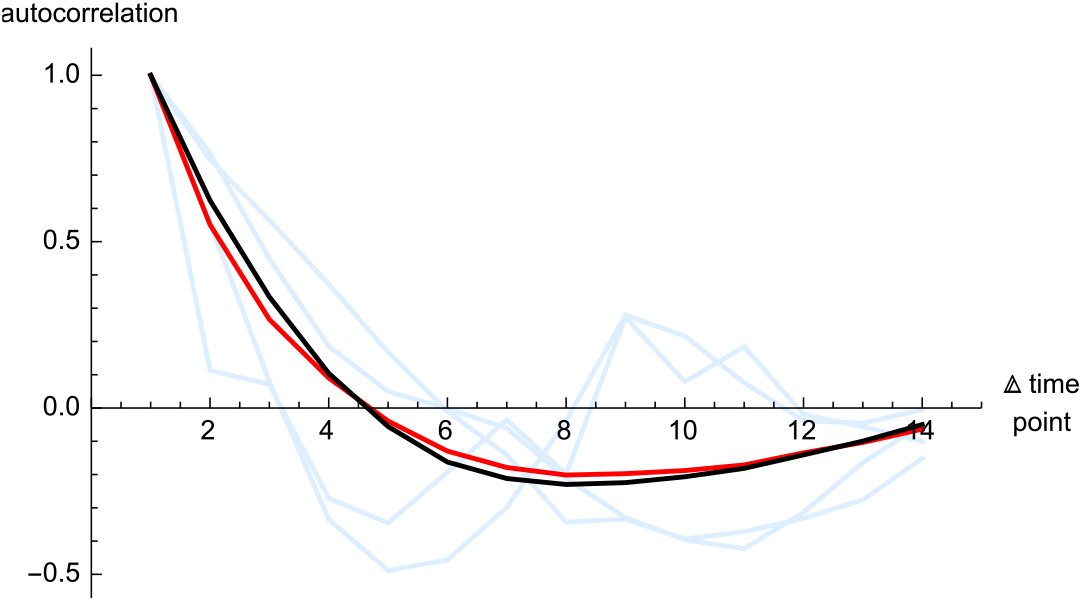

The model agreed very closely with the experimental data when *a* was set to 1, and did not agree for other values of *a*. This implies that the autocorrelation time is infinite, so old noise values are never forgotten.

In retrospect, this makes sense because intracellular YFP and CFP amounts are cumulative, so that any stochastic variation at one time point contributes to the fluorescence at all future time points. The figure shown below investigates this effect further by showing the root mean square deviation of normalized YFP and CFP experimental data away from the values for the same cells at their initial time points. As usual, YFP data are shown in yellow and CFP in cyan, and there is a separate line for each pheromone dose level. Dashed black lines are guides to the eye, showing curves that are proportional to *t*^1/2^.

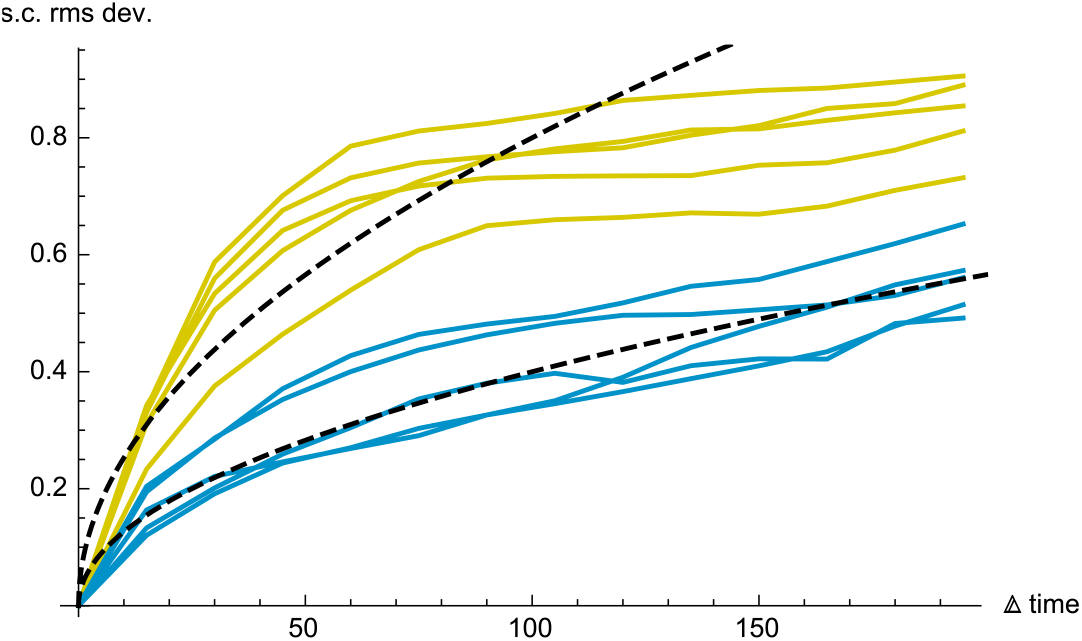

If the stochastic noise were forgotten after time τ, then these curves would be expected to level off exponentially to approach a flat line, also with characteristic time τ. On the other hand, if the stochastic model described above were correct and *a* was equal to 1, then the model responses would be random walks and the standard deviation would increase as *t*^1/2^. In fact, the YFP responses partially level off after about 50 minutes but then continue increasing at a slow but constant rate, implying that the actual situation is somewhere between these limiting cases.

These results have several implications for information transmission. First, taking multiple measurements is still a useful way to reduce the effects of noise, but the information about the response does not grow linearly with the number of measurements. Instead, measurements become less informative over time because the responses include more cumulative noise. Second, the coefficient of variation values that were quantified elsewhere in this work are not fixed quantities but depend weakly on the time duration of the experiment, due to the fact that the rms deviations of the normalized data increase over time. Finally, measurements taken shortly after pheromone addition are likely to be more informative than those taken later, again due to cumulative noise. However, this effect is partially offset by the fact that signals are stronger later on.

## Notes

### Competing Interest Statement

The authors have declared no competing interest.

